# Reproducible detection of antigen-specific T cells and Tregs via standardized and automated activation-induced marker assay workflows

**DOI:** 10.1101/2025.07.15.664847

**Authors:** Torin Halvorson, Gabrielle Boucher, Daniel Yokosawa, Jessica Qing Huang, Rosa Garcia, Lieke Sanderink, Lan Chen, Lorraine Liu, J. Ernesto Fajardo-Despaigne, Jasper Halvorson, Ryan Brinkman, Suzanne Vercauteren, Sylvie Lesage, Jonathan Bramson, John D. Rioux, Sabine Ivison, Megan K. Levings

**Author notes:** Equal contribution (Torin Halvorson, Gabrielle Boucher). Co-senior authors (Sabine Ivison, Megan K. Levings).

## Abstract

Activation-induced marker (AIM) assays are a promising tool to track antigen-specific T cells, but methodological heterogeneity between research groups hinders their clinical utility. To evaluate AIM assay reproducibility, we conducted a multi-site study of SARS-CoV-2 and cytomegalovirus AIMs. We found inherent variability in AIM assays and optimized approaches to enhance reproducibility, including a standardized workflow to minimize technical variability and a generalizable Box-Cox transformation-based statistical method to optimize calculation of AIM stimulation responses. We further standardized AIM data analysis through development of automated flow cytometric gating software, which had superior reproducibility compared to manual analysis. We also characterized antigen-specific Tregs, finding that gating on a combination of CD134, CD137, FOXP3 and HELIOS optimally detects antigen-specific cells. The combined methodology results in a high degree of reproducibility within and between research groups and provides a comprehensive foundation from which standardized AIM assays can be implemented across diverse scientific and clinical settings.

**MOTIVATION:** Reliable detection of antigen-specific T cells is critical to understand immune responses to infection and vaccination, and has translational potential to monitor T cell responses across diverse clinical settings. Activation-induced marker (AIM) assays offer a variety of advantages over methods such as ELISPOT and tetramers, but are limited by methodological heterogeneity between research groups and a lack of standardized protocols. As such, the degree to which AIM assay results are reproducible is unknown. Key variables such as cell source, media, stimulation time, marker selection for CD4^+^ T cells, CD8^+^ T cells and regulatory T cells; as well as data analysis parameters such as flow cytometric gating strategies and mathematical correction for ‘background’ AIM^+^ frequencies in unstimulated control samples, have not been rigorously studied. To address this, we comprehensively characterized variability in AIM assays, including within and between operators and across multiple research centres, and sought to optimize a standard AIM workflow to enhance reproducibility at both the experimental and analytical levels.

## INTRODUCTION

Antigen-specific T cells are critical components of the adaptive immune response to foreign pathogens. Both CD4^+^ and CD8^+^ T cell responses are essential: CD4^+^ conventional T cells (Tconvs) produce lineage-specific cytokines and coordinate activation of innate and adaptive immune cells, while CD8^+^ T cells directly kill infected host cells^1–4^. Upon T cell receptor (TCR)-mediated recognition of cognate antigen along with costimulatory and cytokine signals, CD4^+^ Tconvs and CD8^+^ T cells are activated, proliferate and produce pro-inflammatory cytokines^3,4^.

By contrast, CD4^+^FOXP3^+^ regulatory T cells (Tregs) suppress excessive immune responses and pathological tissue damage, but may also limit immune responses to pathogens^5–7^. Following pathogen clearance, a subset of antigen-specific T cells survive as memory T cells which rapidly respond to subsequent antigen exposure, making them vital for long-lasting immunity^2,8^.

The essential role of T cells in pathogen immunity is exemplified by the COVID-19 pandemic. Robust T cell responses are induced by SARS-CoV-2 infection and/or COVID-19 vaccination, correlate with reduced disease severity and neutralizing antibody production^9–12^, and retain cross-reactivity to antibody-evading variants^13–17^. Indeed, studies in mouse and non-human primate models of SARS-CoV-2 infection demonstrated a critical role for T cells in protection, independently of humoral responses^18,19^.

Beyond SARS-CoV-2, T cell immunity to a wide range of pathogens is critical. For example, human cytomegalovirus (CMV) infects >80% of the global population^20^, and causes a persistent but asymptomatic infection in most individuals^21,22^. CMV-specific T cells are induced by infection and expand throughout life, accounting for up to ∼10% of the memory T cell compartment in older individuals^23^, and, in animal models, are essential for control of viral replication^21,22^. However, in individuals undergoing immune-suppressive therapy, CMV reactivation can cause systemic infection, representing a major cause of mortality after solid organ or hematopoietic stem cell transplantation^24–26^. Deficiency in CMV-specific T cells predicts increased risk of CMV disease in both of these patient populations^25,26^, and early clinical trials have shown benefits of cell therapy with CMV-specific T cells^27–31^.

Clinically-relevant methods to detect antigen-specific T cells are needed to evaluate T cell immunity to infections (including active and latent pathogens), vaccines, and many other applications. In blood, antigen-specific T cells are usually rare^32^, but can be detected via several methods, such as antigen-stimulated proliferation and cytokine production or tetramer staining. However, these assays have several limitations. Proliferation-based assays are time- and resource-intensive, and do not quantify the number of ex vivo T cells^32,33^. Moreover, antigen-non-specific ‘bystander’ activation mediated by cytokine production can confound longer proliferation assays, altering cell phenotypes and causing response overestimation^32,33^. Cytokine-based assays, such as intracellular staining (ICS) and ELISPOT, including interferon-γ release assays (IGRA), can be more specific but do not allow deep cell phenotyping and are limited to cytokine-producing T cells^32–36^. Antigen-specific TCR staining with fluorescently labelled MHC-peptide tetramers is considered the most reliable method to detect and isolate antigen-specific T cells, but requires HLA typing and prior knowledge of TCR epitopes, resulting in limited clinical utility.

Activation-induced marker (AIM) assays have emerged as a practical alternative to quantify antigen-specific T cells. AIM assays take advantage of surface markers upregulated by T cells in response to TCR stimulation, and use flow cytometry to detect combinations of markers specifically expressed by memory, antigen-specific T cells^37^. This approach enables capture of a broad range of antigen-specific T cell responses without requiring prior knowledge of TCR epitopes or their HLA restriction^32,35^. AIM assays also permit the analysis of live cells, enabling downstream cell sorting and further testing of the population of interest^37,38^. Due to their versatility, AIM assays have been used for applications such as detecting rare and/or cytokine-negative antigen-specific T cells^34,37^, evaluating T cell responses to COVID-19 infection^10–12,39,40^ and assessing vaccine immunogenicity^9,15,41–43^.

A plethora of AIMs have been used to identify antigen-specific CD4^+^ or CD8^+^ T cells. In CD4^+^ T cells, CD25 is strongly upregulated upon activation, but is also constitutively expressed on Tregs^32^. CD69 is also expressed in activated CD4^+^ and CD8^+^ T cells, but like CD25, can be susceptible to bystander activation^32^. AIM assay specificity for antigen-specific T cells can be improved by analyzing CD25 or CD69 in combination with a second AIM marker. For example, in combination with CD25, CD134 is a sensitive and specific marker for antigen-specific CD4^+^ T cell activation^34,37,44,45^. CD134 can alternatively be combined with CD274 (PD-L1), reportedly improving detection of Tconvs and excluding Tregs^37^. CD154 (CD40L)^46^ also has utility in detecting CD4^+^ T cell AIM responses. In both CD4^+^ and CD8^+^ T cells, CD137 (4-1BB) is a another marker often used in combination with CD69 or CD134^12,37,40^. CD107a, a marker of cytotoxic degranulation, has been used as an AIM for CD8^+^ T cells^9,42,47^, with its expression correlating with cytotoxicity and IFN-γ production, with similar or greater sensitivity than IFN-γ ICS^42,48^. Combinations of multiple AIMs using Boolean AND/OR gating can increase the sensitivity of AIM assays for antigen-specific T cells^49^.

In recent years, there has been a heightened awareness of the need for transparency and reproducibility in biomedical research^50^. It is therefore critical to develop and implement standard methodological practices for laboratory protocols and assays^51,52^. Despite their potential to monitor immunity to infection and vaccination, clinical application of AIM assays is hindered by a lack of methodological consensus. The choice of culture medium, cell source, stimulation time, AIMs and data analysis pipelines is highly heterogeneous and rigorous reproducibility testing has not been reported. Variability in handling background AIM^+^ signals in unstimulated controls is an additional source of heterogeneity, particularly as low-frequency cell populations are the targets of interest. Thus, here we sought to optimize and standardize AIM assay protocols to detect antigen-specific CD4^+^ Tconvs, Tregs and CD8^+^ T cells, both at the procedural and analytical levels.

We first quantified the variability associated with typical AIM assays and demonstrate that a novel Box-Cox transformation-based method corrects for background unstimulated AIM^+^ frequencies in a way that more appropriately reflects the mathematical relationship between unstimulated and stimulated AIM datasets, reducing technical variability and enhancing signal detection. We further tested key points of protocol variation to determine optimal AIM assay parameters, demonstrating that AIM assays can be performed effectively using cryopreserved peripheral blood mononuclear cells (PBMCs) in serum-free media with a 20-h stimulation period. Inter-lab protocol reproducibility was then tested across four research centres in Canada, and analysis using newly developed automated AIM gating software was compared against manual analysis. We also provide a rigorous characterization of AIM kinetics in Tregs and show that the combination of CD134 and CD137 optimally detects antigen-specific Tregs. Our work provides simple and reproducible protocols for AIM assays that could facilitate their standard implementation as an efficient and sensitive method for clinically evaluating T cell responses to infection and vaccination.

## RESULTS

### AIM assays are inherently variable

A typical AIM assay is performed using whole blood or PBMCs, freshly isolated or cryopreserved, followed by incubation with the antigen of interest (in peptide or whole protein format) and stimulation for 6-48 h^33^. As published AIM assay protocols differ widely, we systematically tested key points of protocol variation across a range of CD4^+^ and CD8^+^ AIMs across diverse conditions as part of the Canadian Autoimmunity Standardization Core (CAN-ASC). We assessed the contribution of serum in culture media, effect of cryopreservation, stimulation time, antigen format and activation markers to overall variability (**Figure 1A**). We employed a 14-colour flow cytometry panel which included 7 AIMs and Treg markers (**Table S1**).

**Figure 1.**
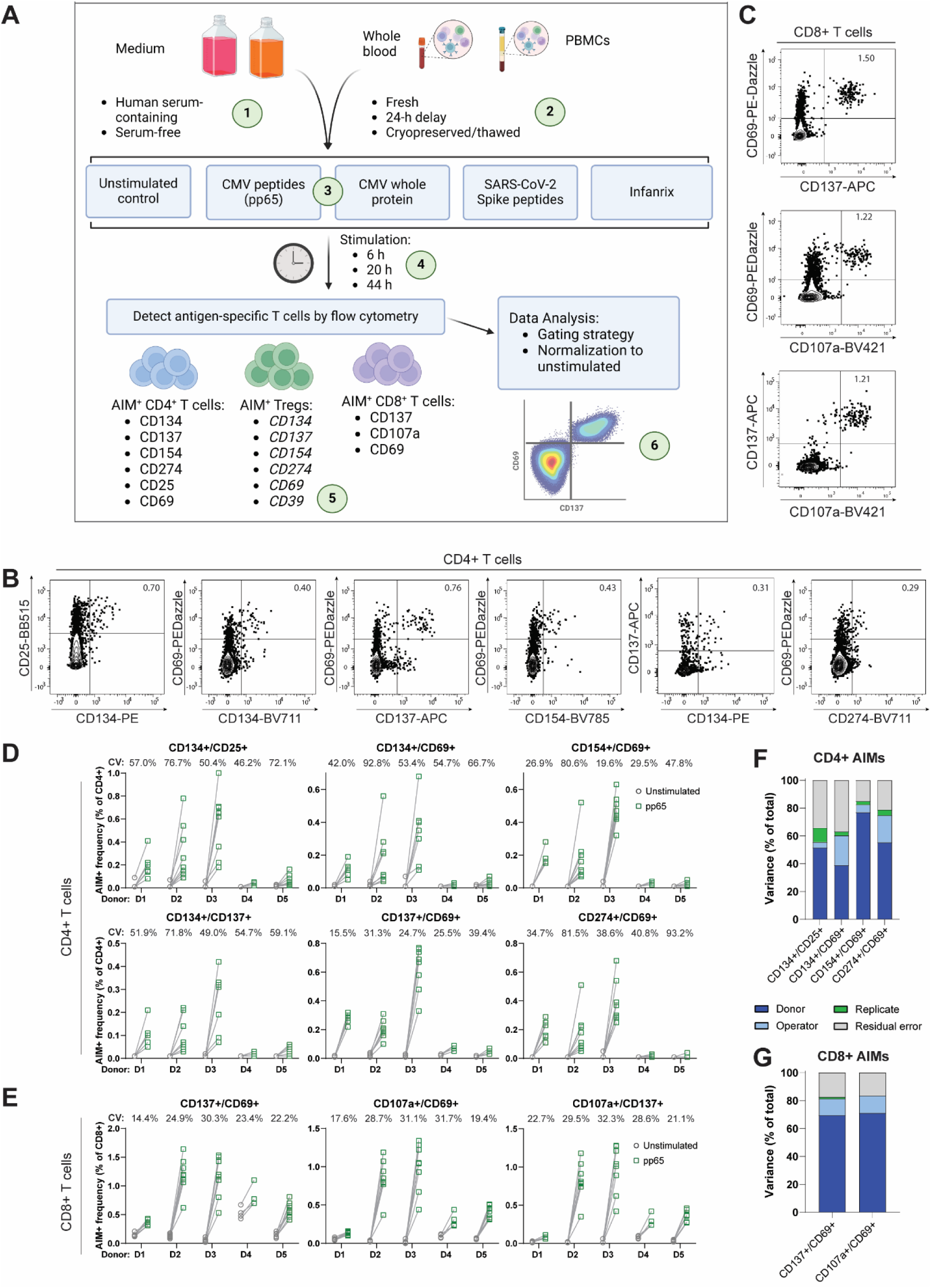
Activation-induced marker assays are inherently variable. (**A**) Schematic depicting the AIM assay procedure and protocol variations that were tested: (1) presence/absence of human serum, (2) use of whole blood, or fresh or frozen PBMCs, (3) stimulation with peptide, whole protein or other antigens; (4), stimulation time; (5) AIM marker combinations and (6) and data analysis strategy. (**B-E**) Replicate AIM assays were performed on cryopreserved PBMCs from 5 donors, with 4-8 technical replicates per donor spread across 2-4 independent experiments. PBMCs were stimulated for 20 h with CMV pp65 peptides and AIM responses were measured by flow cytometry as CD4^+^ or CD8^+^ T cells co-expressing the indicated AIMs. (B and C) Representative dot plots showing AIM^+^ cells after gating on CD4^+^ (B) or CD8^+^ (C) T cells. (D and E) paired raw AIM^+^ frequencies among CD4^+^ (D) and CD8^+^ (E) T cells. Percentage coefficients of variation (CV) are displayed above the responses for each donor. Each line represents the paired unstimulated control and pp65-stimulated AIM signal from a single technical replicate within one donor. (**F and G**) The source(s) of variance contributing to variability in pp65-stimulated CD4^+^ (F) and CD8^+^ (G) AIM^+^ cell frequencies were calculated after subtracting the corresponding unstimulated AIM^+^ frequencies. A mixed effects model with donor, experiment (technical replicate) and operator (the individual performing the AIM assay) as random effects was used. Note that for CD137^+^/CD69^+^ (CD4^+^) and CD107a^+^/CD137^+^ (CD8^+^) data, the mixed effects model failed to converge and variance components could not be calculated. Graphs show variance components expressed as percentages of total variance.

To characterize AIM assay variability, we performed CMV pp65 peptide AIM assays with 4-8 technical replicates of cryopreserved PBMCs from 5 healthy donors (**Table S2**) in at least 2 independent experiments. Antigen-specific CD4^+^ T cells were detected as CD134^+^/CD25^+^, CD134^+^/CD69^+^, CD137^+^/CD69^+^, CD154^+^/CD69^+^, CD134^+^/CD137^+^ or CD274^+^/CD69^+^ and CD8^+^ T cells as CD137^+^/CD69^+^, CD107a^+^/CD69^+^ or CD107a^+^/CD137^+^ (**Figures 1B, 1C and S1**). A coefficient of variation (CV) of <30% between technical replicates has been proposed as an acceptable threshold for clinical standardization of T cell-based assays^53^. AIM^+^ frequencies in stimulated CD4^+^ T cells showed high technical variability, with CVs generally >30%, though CD137/CD69 was a notable exception (**Figure 1D**). By contrast, CD8^+^ AIM assays were more reproducible, with CVs generally <30% (**Figure 1E**).

Analysis of the variance source revealed that for CD4^+^ T cells, only ∼40-75% of assay variance was related to the donor, with a substantial proportion attributable to the replicate (including technical replicates within a single experiment and across independent experiments) and operator (the individual performing the experiment) (**Figure 1F**). In contrast, replicate variance contributed little to CD8^+^ AIM assays, although the operator accounted for ∼10% of the total variance (**Figure 1G**). These data show that AIM assays are subject to technical variability, even when conducted using the same samples and protocol at a single site. In the subsequent experiments and analyses, we sought to reduce AIM assay variability through standardization of the experimental and analytical workflow.

### A Box-Cox transformation-based correction for background reduces AIM assay variability and enhances signal detection

A factor confounding AIM assay analysis is the presence of AIM^+^ T cells in the absence of stimulation, i.e. ‘background’. Researchers variably correct for this background by subtracting or dividing by the corresponding AIM^+^ frequency in the unstimulated control. However, there are no guidelines on how to determine which approach (if either) is appropriate. Subtraction is the most commonly used method^10,34,37,42,49,54^, but is only mathematically correct if the AIM^+^ cells in the unstimulated condition represent random ‘noise’, meaning they are additive to the stimulated AIM signal and have no influence on the response to stimulation. However, if technical or biological factors affect the AIM response, then AIM^+^ frequencies in unstimulated and stimulated conditions may have a non-additive relationship. In this case, division to calculate a ratio of stimulated to unstimulated frequencies is a better expression of stimulation. However, division can lead to substantial variability if unstimulated frequencies are close to zero.

To investigate the potential effects of subtraction versus division, we analyzed the relationship between unstimulated and SARS-CoV-2 Spike peptide-stimulated AIM^+^ frequencies in our previously published cohort of solid organ transplant recipients who received three doses of a COVID-19 mRNA vaccine (n = 33)^55^. In parallel, we applied this analysis to a subset of our current CAN-ASC study cohort of healthy donors (n = 6) stimulated with CMV pp65 peptides in triplicate AIM assays at each of 4 geographically distinct research sites (**Table S2**, described further below). We calculated the stimulated AIM signals for CD4^+^ (CD134/CD25) and CD8^+^ (CD137/CD69) T cells by subtracting the background and found the result significantly and positively correlated with the AIM^+^ frequency in the corresponding unstimulated condition (**Figures 2A and 2B**), indicating that subtraction incompletely removed the effect of background on the net AIM signal. By contrast, division tended to overcorrect, producing trending or significant inverse correlations between unstimulated and stimulated/unstimulated ratios of AIM^+^ frequencies, again suggesting that division failed to eliminate the effect of background on the net AIM signal. These similar observations across two independent datasets led us to conclude that neither subtraction-nor division-based approaches can be universally recommended to correct for background AIM^+^ frequencies in AIM assays.

**Figure 2.**
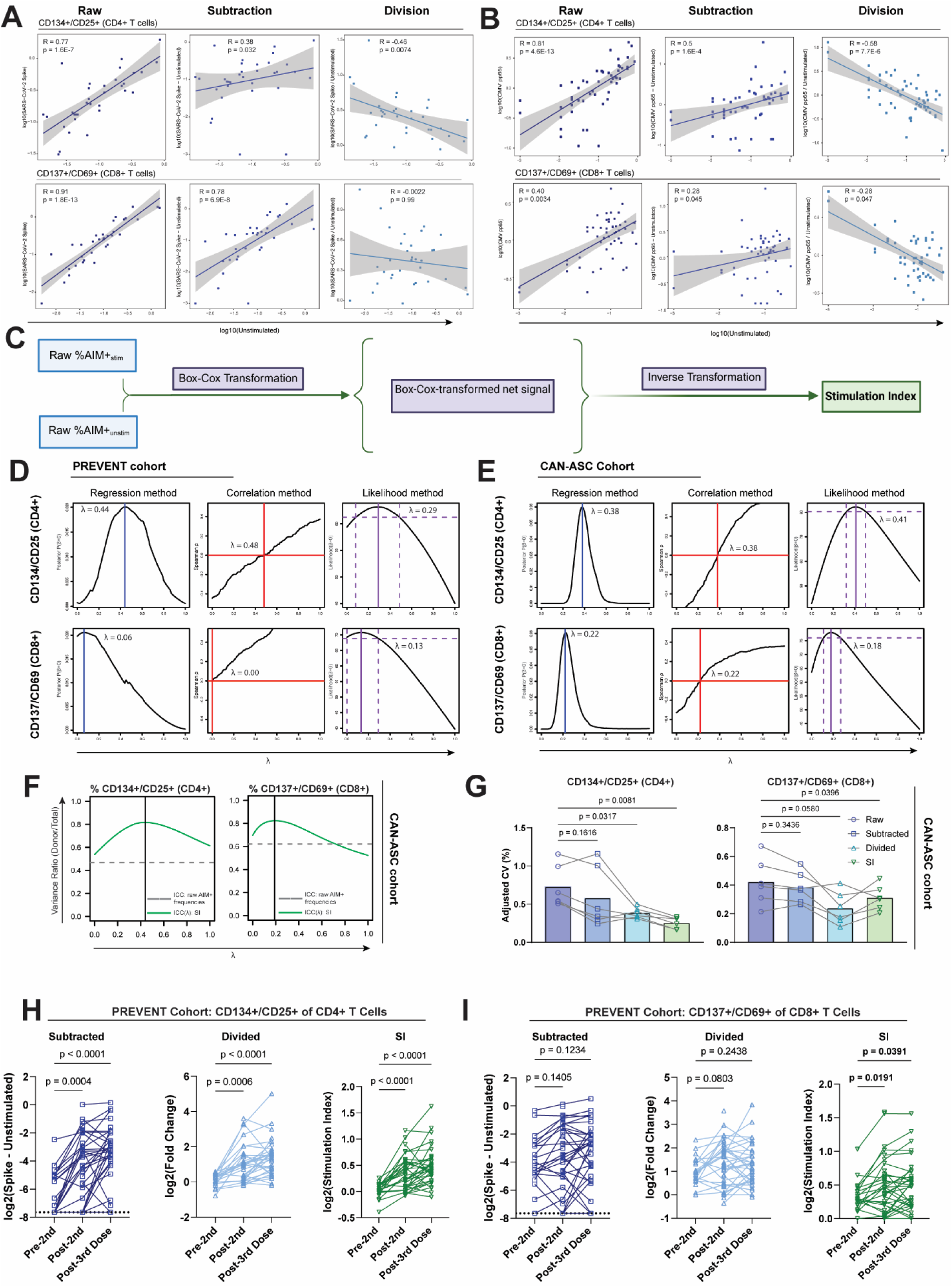
Box-Cox transformation-based correction for background enhances signal detection and reduces variability. (**A**) Re-analysis of previously published AIM assay data from SARS-CoV-2 Spike peptide-stimulated PBMCs from solid organ transplant recipients (n = 33) vaccinated with three doses of a COVID-19 mRNA vaccine (PREVENT cohort)^1^, or (**B**) new analysis of CMV pp65-stimulated PBMCs from 6 healthy donors assayed across 3 independent experiments at each of 4 distinct research centres in the CAN-ASC cohort (n = 52 total). Correlation and linear regression analyses of unstimulated AIM^+^ frequencies with raw antigen-stimulated AIM^+^ frequencies (left panel), or net responses after subtracting (centre panel) or dividing by (right panel) the unstimulated condition. CD134^+^/CD25^+^ frequencies among CD4^+^ T cells (upper panel) and CD137^+^/CD69^+^ frequencies among CD8^+^ T cells (lower panel) are presented. Spearman ρ and p-values are shown from Spearman correlation tests. The solid line is derived from linear regression analysis. For the CAN-ASC cohort, regression and correlation analyses were performed after correcting for between-donor variation (**C**) Schematic showing the workflow of the Box-Cox transformation method of correcting for AIM^+^ cells in the unstimulated control. Raw AIM^+^ frequencies are transformed using the Box-Cox formula with a user-defined parameter l؝[0,1] prior to subtracting to obtain a net signal on the Box-Cox-transformed scale. Inverse Box-Cox transformation of the difference restores the AIM response to the original scale, yielding the stimulation index (SI). (**D and E**) Three statistical methods to estimate optimal values of λ are presented, seeking to minimize correlation between the unstimulated and net stimulated AIM values. Left panel: linear regression, to identify λ giving the highest probability of zero slope (̂β = 0), expressed as a posterior probability distribution for λ. Centre panel: Spearman correlation to identify the value of λ resulting in an estimated zero correlation. Right panel: likelihood profile for λ based on linear regression to estimate the probable optimal value of λ. Values were estimated for the (D) PREVENT or (E) CAN-ASC cohorts. (**F**) Analyses of the CAN-ASC cohort showing ratios of biological (between-donor) variance to total variance (intra-class correlation, ICC; green solid line) plotted as a function of λ using the Box-Cox-corrected stimulation index for CD4^+^ and CD8^+^ AIM responses to CMV pp65. ICC ratios for raw (untransformed) AIM^+^ frequencies are indicated by the grey dotted line. (**G**) Comparison of CVs calculated between technical replicates within individual donors for raw AIM^+^ frequencies, subtraction of or division by the unstimulated AIM^+^ frequency, or Box-Cox-corrected SI. Each point is the CV of technical replicates from one donor, representing within-donor technical variability. P-values are shown from Dunnett’s multiple comparisons test following one-way repeated measures ANOVA. CVs were adjusted to account for the effect of a change in distribution using bootstrap estimates, where the same transformation was repeatedly applied after randomly replacing unstimulated values. (**H and I**) Re-analysis of AIM responses for (H) CD4^+^ and (I) CD8^+^ T cells in n = 40 solid organ transplant recipients from the PREVENT cohort throughout a three-dose COVID-19 mRNA vaccination schedule calculated by subtracting (left) or dividing by (centre) the AIM^+^ frequencies in the unstimulated condition, or Box-Cox-corrected SI values (right). P-values were calculated from log_2_-transformed data using a mixed-effects model with Dunnett’s multiple comparisons test. The bolded p-values denote significant differences between timepoints detected only by analysis of Box-Cox-corrected SI values. See also Figures S2-S5 and Document S1.

To define a more generally applicable approach to isolating the effect of stimulation on AIM signals from background, we leveraged the Box-Cox transformation^56,57^ to analyze AIM stimulation data. The Box-Cox transformation is a family of power transformation known to help in stabilizing variance and render models more linear^56,57^. Transformation of antigen-stimulated and unstimulated AIM frequencies using the Box-Cox transformation prior to subtraction, followed by inverse transformation (**Figure 2C**), is equivalent to subtraction, division or an intermediate operation on the linear scale depending on the value of a user-defined parameter lambda (l؝[0,1]). Using this method, we calculated an AIM stimulation index (SI) in three steps:

1. Apply the transformation: 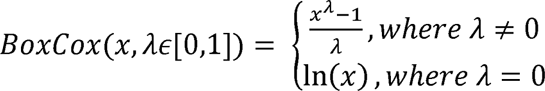
2. Obtain the net stimulated value using subtraction: *β = BoxCox(%AIM^+^_stim_*) – *BoxCox(%AIM^+^_unstim_*)
3. Return to original scale: 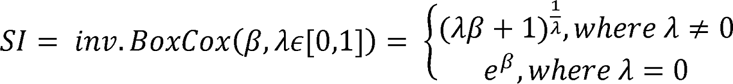

When λ = 0, the SI is equivalent to division by the unstimulated AIM^+^ frequency (fold change); when λ = 1, the result is a linear subtraction plus 1. When λ is between 0 and 1, the function is intermediate between division and linear subtraction. This avoids the pitfall of dividing by very low unstimulated values, while still controlling for some multiplicative effects. The method is presented in further detail in **Document S1,** including additional modifications to the formula to remove the translation at λ = 1, defining a modified (m)SI.

We next sought an optimal λ value which would minimize the residual correlation or linear relationship between the net stimulated and unstimulated AIM value. We employed three complementary mathematical methods to estimate the optimal λ-value, using data from both the PREVENT and CAN-ASC cohorts (see **Document S1** and methods). In both cohorts, we found that the optimal λ-value often fell far from the extremes of division (λ = 0) and subtraction (λ = 1), with results depending on the specific AIMs (**Figures 2D, 2E and S2A-D**). We next re-analyzed the relationship between unstimulated (background) AIM^+^ frequencies and net AIM Box-Cox-corrected signal in the PREVENT and CAN-ASC datasets. Here, for simplicity and consistency, we selected λ = 0.5 as a mathematically intermediate operation between division and subtraction. For most AIMs and in both cohorts, accounting for background AIM^+^ frequencies with the Box-Cox correction method resulted in weak or negligible correlations between unstimulated and net AIM signals observed with subtraction of, or division by, unstimulated AIM^+^ frequencies (**Figures S2E and S2F)**.

As a measure of variability, we used the intra-class correlation value (ICC). ICCs >0.5 indicate biological and experimental variation are equal and an ICC approaching 1 indicates sensitive signal detection, while ICC <0.5 indicates that experimental error will obscure biological differences. Using the multi-centre CAN-ASC dataset, we found that ICCs were consistently higher when using a Box-Cox-corrected SI with intermediate λ values, when compared with subtraction of (λ=1) or division by (λ=0) the unstimulated control, or with the raw data (**Figures 2F, S3A and S3B**). Similar results were obtained using the scale-independent mSI (**Figures S3C and S3D)**.

Given the desirable increase in ICCs with Box-Cox-corrected SIs, we next investigated whether this method reduced CVs between technical replicates. The CV is distribution-dependent, and not robust to most data transformation, including translation. To account for this, we employed a bootstrapping method to adjust the CVs for the expected impact of the transformation under the hypothesis of uninformative unstimulated AIM+ frequencies. Consistent with the ICC data, Box-Cox-corrected SIs produced lower intra-donor adjusted CVs for key CD4^+^ and CD8^+^ AIMs (**Figures 2G, S4A and S4B**), further supporting the ability of this method to reduce technical variability and enhance signal detection.

We next applied the Box-Cox correction with λ = 0.5 to re-analyze key results from the PREVENT dataset^55^, hypothesizing that reducing technical variability through improved correction for unstimulated AIM^+^ frequencies could enhance detection of differences between vaccination timepoints. While Box-Cox correction did not impair detection of already highly significant differences between CD4^+^ T cell responses at pre-2^nd^ dose, post-2^nd^ dose and post-3^rd^ dose timepoints (**Figure 2H**), it enhanced signal detection among variable CD8^+^ T cell responses, leading to increased detection of significant differences between groups (**Figure 2I)**.

We also asked whether the Box-Cox correction (with λ = 0.5) could be applied together with the 6xAIM method recently reported by Lemieux *et al*^49^ which uses Boolean OR gating to sum cells expressing any of CD134/CD69, CD137/CD69, CD154/CD69, CD134/CD154, CD134/CD137 or CD137/CD154 to detect CMV-specific T cells in a subset of healthy donors from the CAN-ASC cohort (n = 9). Although the 6xAIM method increased the background signal in unstimulated samples, it enhanced detection of CMV-specific CD4^+^ T cells compared with CD134/CD154 or CD134/CD69 (**Figure S5A**), but not with CD134/CD25. Adding CD134/CD25 or CD274/CD69 to the 6xAIM led to trending or significantly improved signal detection. In line with Lemieux *et al.*, the 6xAIM did not enhance AIM signal detection in CD8^+^ T cells compared to CD137/CD69 alone^49^, and we found no benefit of adding CD107a/CD69, CD107a/CD137 and/or CD274/CD69 to the 6xAIM combination (**Figure S5B**).

Taken together, these results indicate that Box-Cox transformation and subtraction using a value of λ between 0 and 1 yields a mathematically intuitive AIM SI that can reduce the effects of technical variability and enhance signal detection. For simplicity, we elected to use λ = 0.5 for all subsequent analyses, but we note that the Box-Cox correction can be customized based on the mathematical properties of a given dataset (see **Document S1**). To facilitate method deployment, we developed the Box-Cox Correction Application^58^ (see Methods) a flexible web-based tool that allows researchers to rapidly generate Box-Cox-corrected SI data from raw AIM^+^ frequencies with no statistical or coding knowledge required.

### CD4^+^ and CD8^+^ AIM responses are reliable in serum-free media and various cell sources

Media containing human serum is commonly used in AIM assays but batch-to-batch variability of this reagent limits standardization. To test AIM responses in serum free media, we stimulated PBMCs with CMV pp65 peptide in X-Vivo + 10% human serum (XH), serum-free X-Vivo (XV), or serum-free ImmunoCult-XF (IC). Viability was unaffected by the choice of medium (**Figure S6A**). The geometric mean fluorescence (MFI) of CD4 (but not CD8) was lower in XV and trended lower in IC compared to XH, but separation of CD4^+^ and CD4^-^ populations was maintained (**Figures S6B and S6C**).

We found that although serum-free media caused trending or significant increases in background CD134^+^/CD25^+^ AIM frequencies, similar increases were observed for antigen-stimulated frequencies (**Figure S6D**), thus, no differences in Box-Cox-corrected SIs for CD4^+^ or CD8^+^ T cells were identified (**Figure S6E)**. Importantly, serum-free media did not significantly increase AIM assay variability between technical replicates (**Figure S6F**). Thus, AIM assays can be performed in serum-free media without significantly impacting signal detection.

AIM assays using fresh whole blood are the gold standard for antigen-specific T cell detection, but this cell source has complex logistics. Use of cryopreserved PBMCs enables batch analyses, minimizing experimental variation, but the processes of PBMC isolation and cryopreservation can themselves introduce variability, especially if sample processing is delayed. Using 6 healthy donors, we compared AIM detection in fresh whole blood, whole blood after a 24 h delay, or in PBMCs isolated immediately or 24h after blood draw, and that were or were not cryopreserved. As whole protein antigens require proteolytic processing to generate peptides prior to presentation on MHC molecules, AIM responses could differ from responses to peptide pools depending on the differential antigen processing capacities of the cells present. We therefore compared AIM responses to CMV pp65 peptides and whole pp65 protein across cell sources. In parallel, to extend our findings to more diverse antigen types across different cell sources, we also tested AIM responses to the Infanrix hexa combination vaccine, a component of standard childhood immunization regimens that contains a combination of whole inactivated virus, protein, toxoid and conjugated polysaccharide antigens derived from six bacterial and viral pathogens.

Low cell viability (<60-70%) significantly biases results of PBMC-based assays^52^. We confirmed that although T cell viability was highest in whole blood or freshly isolated PBMCs, it remained >75% in samples processed after 24 h or post-cryopreservation (**Figure 3A**). To compare the reliability of AIM response detection across cell sources, we deemed each AIM signal significantly detectable when the 95% confidence interval of the log_2_-transformed SI did not overlap with 0. CD4^+^ T cell AIMs, and CD137^+^/CD69^+^ responses in CD8^+^ T cells, were generally more reliable in fresh or 24h-delayed PBMCs than in whole blood (**Figures 3B-E**). However, AIM pairs involving CD107a were more consistently detectable in whole blood. A 24-h delay in whole blood AIM assay initiation consistently resulted in loss of detection of weaker CD4^+^ and CD8^+^ responses, such as responses to Infanrix for several AIMs. CD4^+^ and CD8^+^ AIM signals were reduced in cryopreserved PBMCs but remained significantly detectable, particularly in CMV-stimulated samples. However, a 24 h delay in PBMC processing led to a loss of signal detection in cryopreserved PBMCs with some AIMs, highlighting the necessity of immediate PBMC processing for optimal sensitivity.

**Figure 3.**
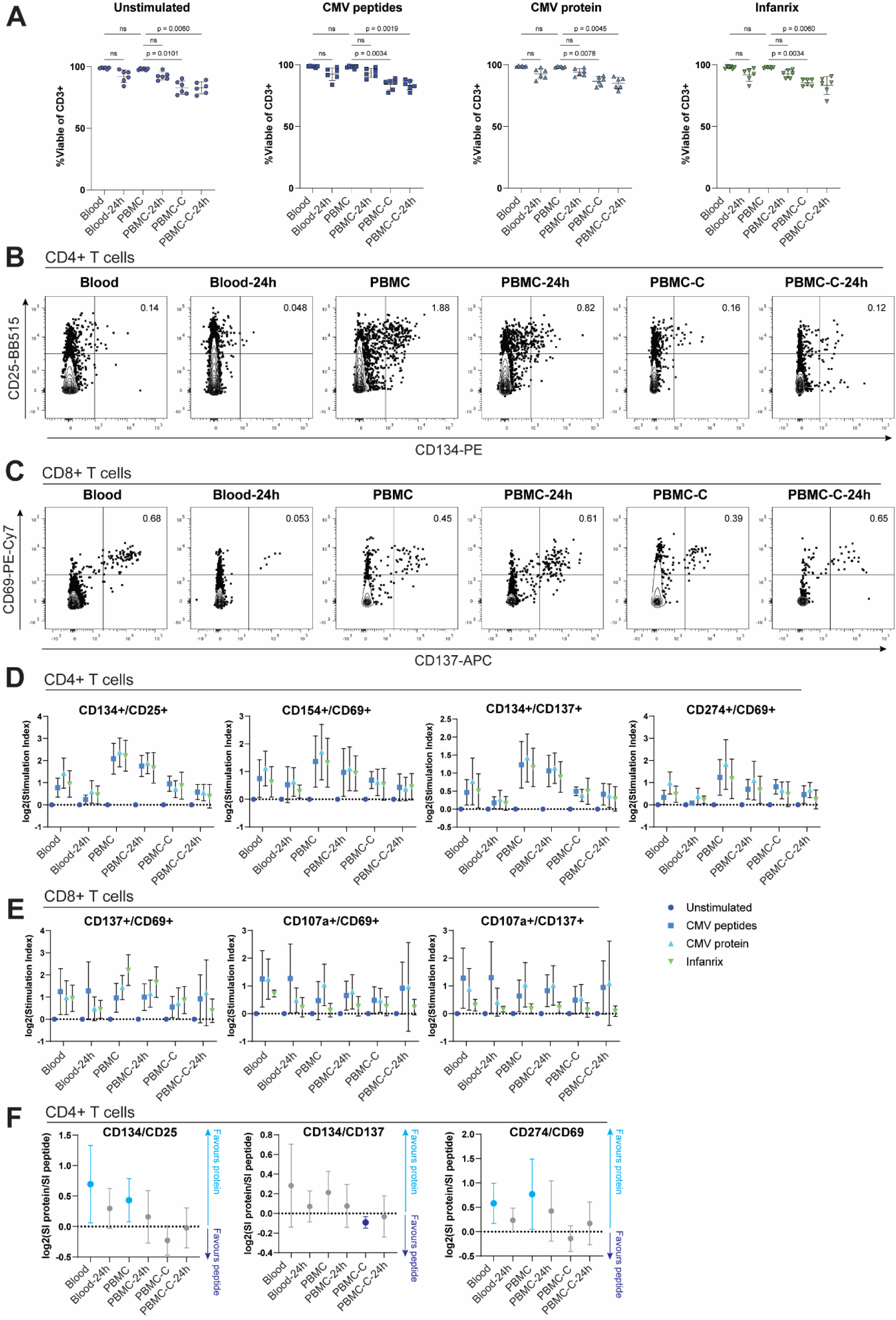
Activation-induced marker (AIM) responses are reliably detectable across various cell sources. Comparisons of AIM responses using fresh whole blood, or fresh (PBMC) or cryopreserved (PBMC-C) peripheral blood mononuclear cells, with or without a 24-h delay in processing (- 24h). Cells were rested overnight and stimulated for 20 h with cytomegalovirus (CMV) pp65 peptides, CMV pp65 whole protein, or Infanrix vaccine. (**A**) Percentage viable (fixable viability dye-eFluor 780-negative) cells of total CD3^+^ T cells measured by flow cytometry in AIM assays. Error bars represent the standard deviation of the mean. P-values are shown for planned comparisons using Dunn’s multiple comparisons test following a Friedman test. (**B-E**) Representative flow cytometry (B and C) and quantification of stimulation index for each AIM relative to the unstimulated control (D and E) among CD4^+^ (B and D) and CD8^+^ (C and E) T cells from each cell source. (**F**) Ratio of AIM stimulation indices between CMV pp65 protein- and peptide-stimulated CD4^+^ T cells in relation to cell source. Positive or negative ratios indicate greater detection of AIM responses with whole protein or peptide stimulation, respectively, while confidence intervals overlapping zero indicate no significant difference between protein and peptide stimulation. (D, E and F) Error bars are 95% confidence intervals calculated on a log_2_ scale. (A-F) Data from n=6 healthy donors. See also Figure S7.

Analysis of AIMs after stimulation with whole CMV pp65 protein compared to peptides showed intriguing trends in combination with cell source. Among CD4^+^ T cells, CD134/CD25, CD134/CD137 and CD274/CD69 AIM signal detection favoured whole protein stimulation when the assay was performed with whole blood or fresh PBMCs, while peptide stimulation was optimal when using cryopreserved/thawed PBMCs (**Figure 3F**). These trends were less clear for other CD4^+^ or CD8^+^ T cell AIM responses (**Figures S7A and S7B**).

### A 20-h stimulation period optimally detects a range of CD4^+^ and CD8^+^ AIMs

AIM assays are performed using stimulation periods ranging from 6-48 h. However, the kinetics of individual AIMs vary, and a ‘consensus’ timepoint where many AIMs can be simultaneously detected has not been rigorously characterized. CD69, CD154 and CD107 are rapidly (2-6 hours) upregulated following TCR stimulation and remain detectable for at least 24 h^9,38,42,47,48,59–61^. In contrast, CD137 and CD274 are optimally detected following longer stimulation periods of 18-24 h^34,37,62–65^, and CD25 and CD134 reach maximum expression after 48 h^33,66,67^. Moreover, kinetics may also be different in whole blood compared with PBMCs.

We thus systematically compared AIM SIs following 6 h, 20 h or 44 h of stimulation with pp65 peptides or whole protein, using whole blood or cryopreserved PBMCs. AIM readouts involving CD134, CD137 and CD274 required stimulation for at least 20 h (**Figures 4A, 4B and S8A**), with optimal timing differing between blood and PBMCs, particularly for CD274/CD69 which were optimally detected at 20 h in blood but at 44 h in PBMCs (**Figures 4B and S8A**). Although some donors displayed strong CD154/CD69 responses in CD4^+^ T cells at 6 h, signal detection was most reliable at 20 h in blood and at 20 or 44 h in PBMCs (**Figures 4B and S8A**).

**Figure 4.**
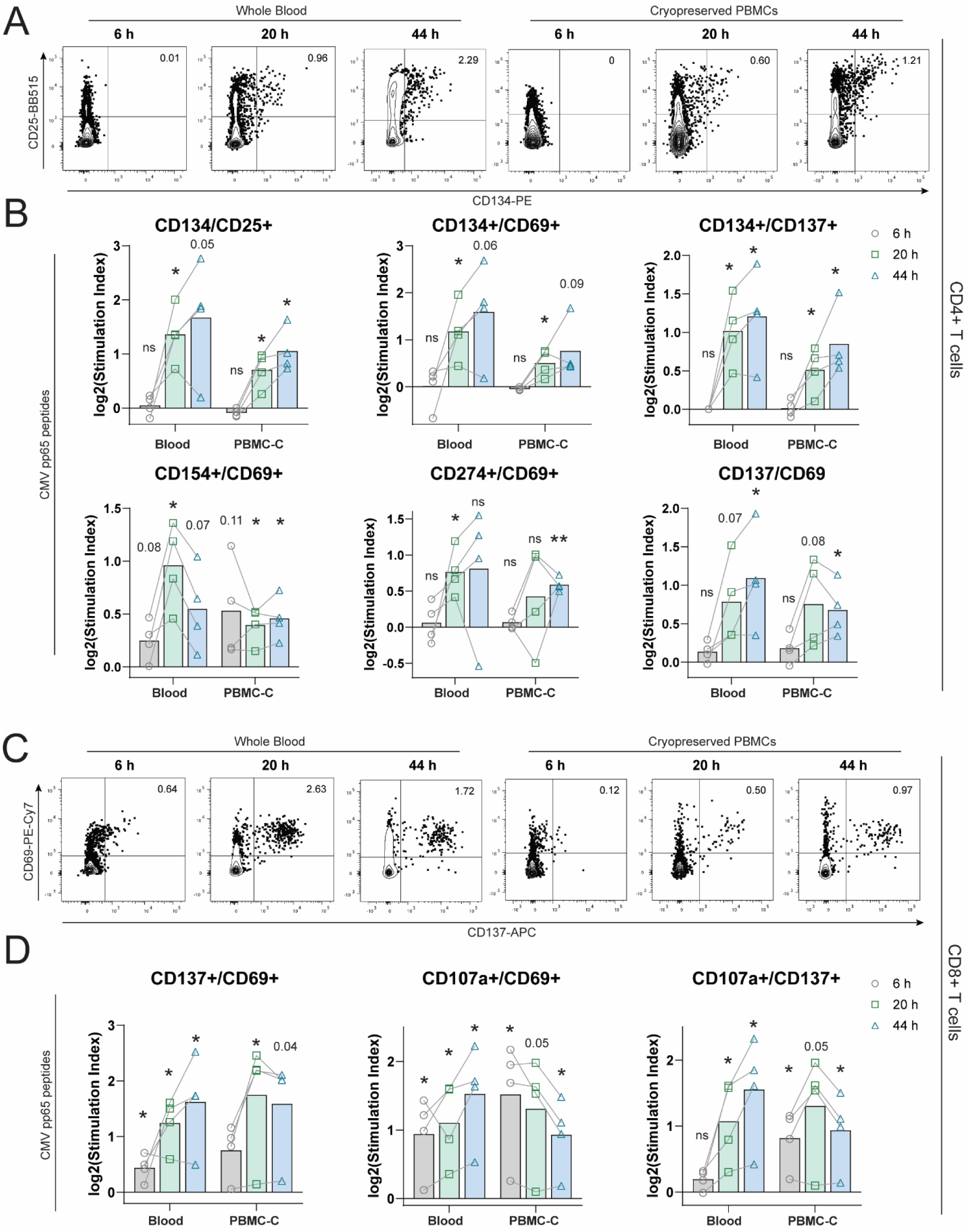
A 20-h stimulation period reliably detects most CD4^+^ and CD8^+^ T cell activation-induced markers (AIMs). (**A**) Representative and (**B**) quantified flow cytometry data showing CD134^+^/CD25^+^ cells among CD4^+^ T cells following stimulation of cryopreserved PBMCs (PBMC-C) or fresh whole blood with CMV pp65 peptides for 6, 20 or 44 h. (**C**) Representative and (**D**) quantified flow cytometry showing CD137^+^/CD69^+^ cells among CD8^+^ T cells following stimulation of PBMC-C or fresh whole blood with CMV pp65 peptides for 6, 20 or 44 h. (B and D) SIs are represented as log_2_- transformed data. P-values are shown from one-sample t-tests, with AIM signals considered present when the mean log_2_-transformed SI significantly differs from zero. All data are paired samples from n = 4 healthy donors. ns, not significant (p > 0.05); *, p ≤ 0.05; **, p ≤ 0.01; ***, p ≤ 0.001; ****, p ≤ 0.0001. See also Figure S8.

CD8^+^ T cell CD137/CD69 responses were detected at 20 and 44 h, with significant but low-magnitude responses observed at 6 h for peptide stimulation in blood (**Figures 4C, 4D and S8B**). In contrast, AIM pairs containing CD107a yielded significant 6-h responses only in PBMCs, whereas detection in blood was most reliable at 20 h (**Figures 4D and S8B**). Overall, the 20 h stimulation period resulted in significant detection of AIM signals from the broadest range of AIMs, including pairs involving CD25, CD69, CD134, CD137, CD154 and CD107a. Hence, we selected a 20 h timepoint as a consensus for subsequent CD4^+^ and CD8^+^ AIM assays.

### CD134 and CD137 optimally detect antigen-specific CD4+ Tregs

Although AIM assays are commonly used to detect T cells that contribute to protective immunity, antigen-specific Tregs are also induced by infection and vaccination^37,68,69^. While bulk Treg frequencies are increased in severe COVID-19^70–72^, SARS-CoV-2-specific Treg responses are associated with milder disease^73^. CMV-specific Tregs have also been described, comprising a large subset of CMV-responsive CD4^+^ T cells^74^, and are associated with recurrent CMV disease in kidney transplant recipients^75^. Therefore, distinguishing between antigen-specific CD4^+^ Tconvs and Tregs in AIM assays may be clinically important across diverse disease contexts, but Treg AIM markers and their kinetics are not well-characterized.

An important concern is that Treg-associated marker FOXP3 is also transiently expressed by activated Tconvs following TCR stimulation *in vitro*^76^, potentially limiting its utility as a Treg lineage marker. We thus hypothesized that HELIOS, a transcription factor characteristic of stable thymus-derived Tregs^77^, may be a more reliable marker to identify Tregs in AIM assays. We first evaluated the kinetics of FOXP3 and HELIOS expression in CD4^+^ T cells (**Figures 5A and 5B**), finding that mean FOXP3^+^ frequencies among CD4^+^ T cells significantly increased from 6 h (7.1%) to 20 h (9.1%; p<0.05) and 44 h (11.7%; p<0.01) post-stimulation with pp65 peptides. Conversely, FOXP3^+^/HELIOS^+^ double-positive cell frequencies remained similar (∼3-5%) in unstimulated and pp65-stimulated conditions across all timepoints, suggesting that this population represents true Tregs. We therefore gated on FOXP3^+^/HELIOS^+^ double-positive Tregs for subsequent analyses (**Figure S9**).

**Figure 5.**
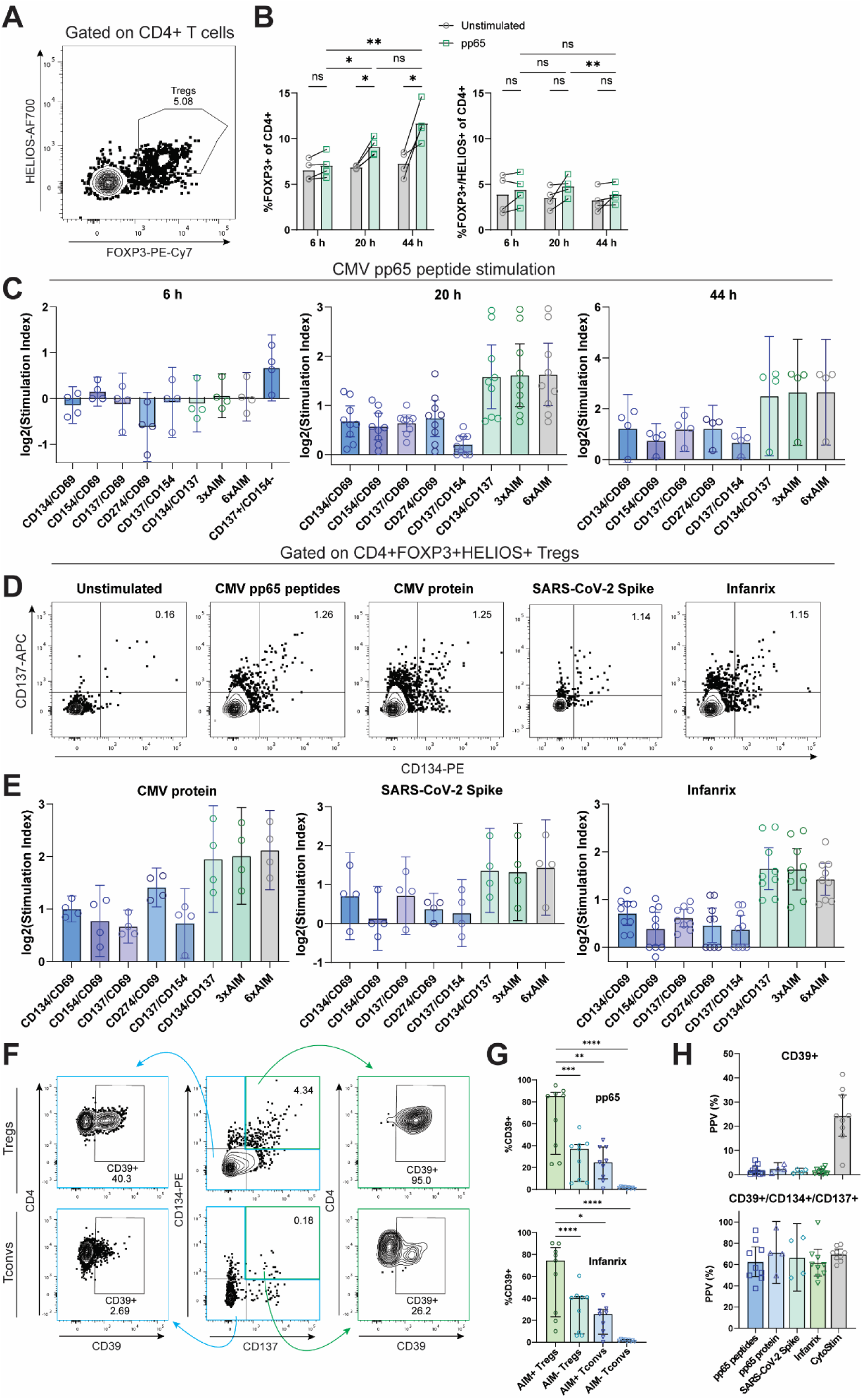
CD134 and CD137 optimally detect antigen-specific regulatory T cells (Tregs). (A) Representative flow cytometric gating strategy for FOXP3^+^/HELIOS^+^ Tregs. (**B**) Effect of stimulation with cytomegalovirus (CMV) pp65 peptides on FOXP3^+^ and FOXP3^+^/HELIOS^+^ cell frequencies among CD4^+^ T cells over time. P-values represent Tukey’s multiple comparisons test following a two-way ANOVA.(**C-E**) Stimulation indices (C and E) or representative flow cytometry (D) for AIM pairs and Boolean combinations in FOXP3^+^HELIOS^+^ Tregs following stimulation of cryopreserved PBMCs with pp65 peptides, CMV pp65 whole protein, SARS-CoV-2 Spike peptides or the Infanrix combination vaccine for 20 h. Boolean AND/OR gating combinations comprise CD134/CD69, CD134/CD137 and CD137/CD69 (3xAIM); or 3xAIM plus CD154/CD69, CD134/CD154 and CD137/CD154 (6xAIM). Stimulation index was determined by normalizing pp65 peptide-stimulated AIM^+^ frequencies to the unstimulated control using the box-cox correction method described in Figure 2. The mean log_2_-transformed stimulation index is shown for each AIM. Error bars represent 95% confidence intervals of the mean. Data represent n = 9 (pp65 and Infanrix 20-h stimulations) and n = 4 (all others) healthy donors. (**F**) Flow cytometric gating strategy to evaluate CD39^+^ frequencies among CD134^+^/CD137^+^ and non-CD134+/CD137+ (comprising CD134^+^/CD137^-^, CD134^-^/CD137^-^ and CD134^-^/CD137^+^ cells) FOXP3^+^/HELIOS^+^ Tregs and FOXP3/HELIOS^-^ Tconvs. (**G**) Median percentages of CD39^+^ events among CD134^+^/CD137^+^ or CD134/CD137^-^ CD4^+^/FOXP3^+^/HELIOS^+^ Tregs or conventional FOXP3/HELIOS^-^ T cells (Tconvs). (**H**) Positive predictive value (PPV) of CD39 (top) or CD39, CD134 and CD137 (bottom) as markers for antigen-specific Tregs (CD134^+^/CD137^+^/FOXP3^+^/HELIOS^+^) among total CD4^+^ T cells. ns, not significant (p > 0.05); *, p ≤ 0.05; **, p ≤ 0.01; ***, p ≤ 0.001; ****, p ≤ 0.0001. See also Figure S9.

Antigen-stimulated Tregs upregulate CD137 earlier than Tconvs (within 5-7 h), but have weak and delayed expression of CD154^64,78,79^ so can be identified as CD4^+^CD137^+^CD154^-^ T cells at 6 h^79,80^. We confirmed these results in FOXP3^+^HELIOS^+^ Tregs, demonstrating CD137^+^/CD154^-^ responses to pp65 peptides at 6 h in Tregs from 3 of 4 donors, with little expression of this combination in Tconvs (**Figure S10A)**. We then systematically compared pp65-specific FOXP3^+^HELIOS^+^ Treg responses across multiple AIM combinations at 6, 20 and 44 h, again considering responses significant if the 95% CI of the log_2_-transformed SI did not overlap with zero (**Figure 5C**). CD25 was not included, as this marker is constitutively expressed by Tregs. At 6 h, no AIM pair was significantly detected in Tregs, but CD134/CD69, CD154/CD69, CD137/CD69, CD274/CD69, CD137/CD154 and CD134/CD137 responses were significantly detectable at 20 h. The combination of CD134 and CD137 yielded the highest SI in Tregs, and the signal strength was not further increased with Boolean combinations of CD134/CD137, CD134/CD69 and CD137/CD69 (‘3xAIM’) or the 6xAIM^49^ (**Figure 5C**). The CD134/CD137 pair was consistently upregulated in antigen-specific FOXP3^+^HELIOS^+^ Tregs following stimulation with pp65 whole protein, SARS-CoV-2 Spike peptides and Infanrix, demonstrating the robustness of this method across diverse antigens (**Figures 5D and 5E**).

The ectonuclease CD39 is expressed on a subset of FOXP3^+^ Tregs in AIM assays^37,81,82^. In 20-h AIM assays, CD39 was expressed in ∼5% of antigen-stimulated CD4^+^ T cells (**Figure S10B**), and generally less than 50% of these cells were FOXP3^+^HELIOS^+^ Tregs (**Figure S10C**). However, CD39^+^ cells were enriched in activated CD134^+^CD137^+^ FOXP3^+^HELIOS^+^ Tregs, compared to non-activated Tregs, or Tconvs (regardless of activation status) (**Figures 5F and 5G**). Overall, CD39 alone yielded variable sensitivity (**Figure S10D**) and low positive predictive value (PPV) for the identification of antigen-specific Tregs, but when combined with CD134 and CD137, the mean PPV increased to >60% across a range of antigenic stimulants (**Figure 5H**). Overall, in line with previous observations in FOXP3^+^ Tregs^81,82^, CD39 is highly associated with activated, antigen-specific stable FOXP3^+^/HELIOS^+^ Tregs and may be useful in combination with other AIM markers if intracellular FOXP3/HELIOS staining is undesirable.

### CMV-specific and SARS-CoV-2-specific AIM assays are reproducible within and between laboratories

To test the reproducibility of our AIM assay protocol, we next conducted a multi-site testing experiment across four centres: British Columbia Children’s Hospital (clinical laboratory), the Human Immune Testing suite at McMaster University, and research laboratories at the Universities of Montreal and British Columbia (**Figure 6A**). Two sites used CytoFLEX cytometers (Beckman Coulter) while two used LSRFortessa cytometers (BD Biosciences); each cytometer type was standardized to the same set of target regions; however, the two types of cytometers were not standardized to each other due to different technologies and scaling. For these tests, we selected an optimized AIM protocol based on our results using a simplified flow cytometry panel (**Table S3**), selecting the AIMs found to be most consistent in preceding experiments (**Figure S11**). PBMC aliquots from healthy donors (n = 6) were cryopreserved without processing delays. PBMCs were thawed, rested overnight and stimulated in IC medium with pp65 or SARS-CoV-2 Spike peptides. After 20 h, antigen-specific T cell responses were detected by flow cytometry and stimulation indices calculated using the Box-Cox correction. Each assay was repeated three times per site (experimental replicates), with each replicate from a different batch of thawed PBMCs, performed on a different day by the same operator (**Figure 6A**). In line with efforts to standardize tetramer-based detection of antigen-specific T cells^53^, a CV of 30% was set as a threshold to define reproducibility.

**Figure 6.**
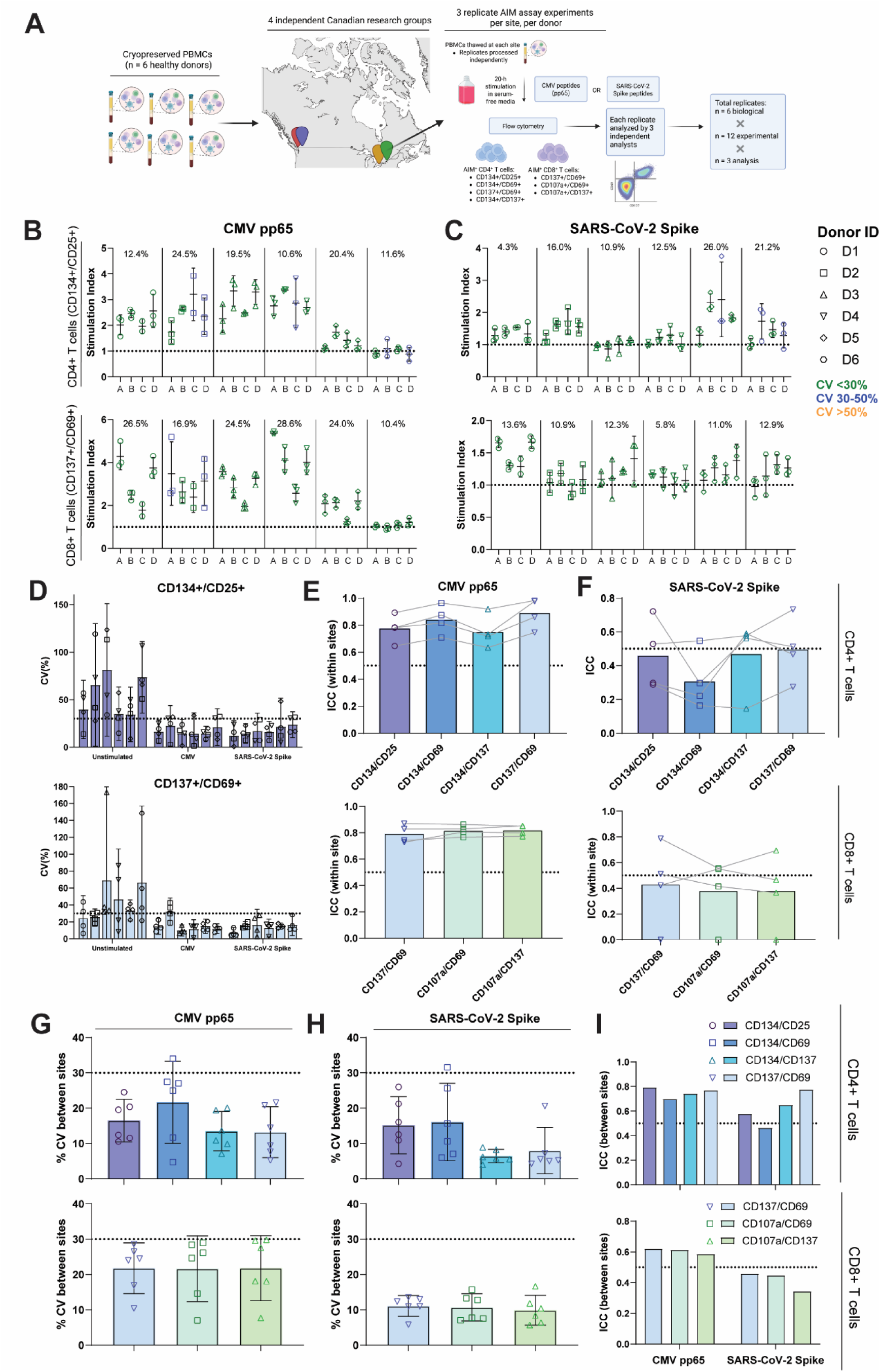
Cytomegalovirus (CMV)- and SARS-CoV-2-specific AIM assays are reproducible within and between geographically distinct research centres. (**A**) Experimental design for AIM assay inter-site testing. Replicate aliquots of PBMCs from n = 6 healthy donors were assayed in three independent experiments (individual points) by four distinct Canadian research groups (coded A-D). Replicate AIM assay experiments (n = 3 per donor, per site) were performed by stimulating PBMCs (thawed independently for each replicate) for 20 h with CMV pp65 or SARS-CoV-2 Spike peptides. (**B and C**) Stimulation indices of CD134^+^/CD25^+^ frequencies among CD4^+^ T cells (top) and CD137^+^/CD69^+^ frequencies among CD8^+^ T cells (bottom). Figure subpanels separate individual donors. Stimulation index was determined by normalizing pp65 peptide-stimulated AIM^+^ frequencies to the unstimulated control using the Box-Cox correction method described in Figure 2. The means and standard deviations of log_2_-transformed stimulation indices are shown for each AIM. Symbol colours indicate the percent coefficient of variation (CV) within each set of three replicates at each site. The overall CV between sites is shown at the top of the panel for each donor. (**D**) Percent CV in CD4^+^ (top) and CD8^+^ (bottom) T cell AIM stimulation indices calculated between technical replicate AIM assays within each site for unstimulated, pp65- stimulated and Spike-stimulated conditions. Each point represents the CV of three technical replicates within one site; bars represent individual donors. CV means with standard deviation are shown. The dotted line represents CV = 30%, a common threshold for acceptable assay reproducibility. (**E and F**) ICCs comparing the technical variability among CD4^+^ (top) and CD8^+^ (bottom) AIM responses to (D) CMV pp65 and (E) SARS-CoV-2 Spike using data from replicate AIM assays from different sites and donors. Bars represents the ICC for each AIM as a mean across all sites. The dotted line indicates ICC = 0.5, the point at which biological and technical variability are equal. (**G and H**) Percent CV in CD4^+^ (top) and CD8^+^ T cell (bottom) AIM responses to CMV pp65 (F) and SARS-CoV-2 Spike (G) between sites, calculated after averaging three technical replicate AIM assays at each site. Each point represents one donor. Error bars are 95% confidence intervals of the mean. The dotted line represents CV = 30%, a common threshold for acceptable assay reproducibility. (**I**) ICCs comparing the calculated technical variability for replicate AIM assays between sites to biological variability between donors for CD4^+^ (top) and CD8^+^ (bottom). The dotted line indicates ICC = 0.5, the point at which biological and technical variability are equal. See also Figures S10-S14.

AIM responses to CMV and SARS-CoV-2 were detected at similar magnitudes between experimental replicates at each site, with CVs <30% for nearly all donor-site combinations for CD4^+^ (CD134^+^/CD25^+^) and CD8^+^ (CD137^+^/CD69^+^) (**Figures 6B-D).** We observed similarly low intra-site variability for all other AIM pairs (**Figures S12A-D, S13A and S13B**), despite high variability in background unstimulated AIM^+^ frequencies, highlighting the importance of appropriate background correction. ICC analysis comparing biological and technical variability at each site demonstrated ICCs consistently >50% for CMV AIM assays (**Figure 6E**), but lower ICCs for SARS-CoV-2 assays (**Figure 6F**). CMV AIM assay reproducibility varied between different markers, with CVs significantly <30% for CD134^+^/CD25^+^, CD134^+^/CD137^+^ and CD137^+^/CD69^+^ in CD4^+^ T cells, and CD137^+^/CD69^+^ in CD8^+^ T cells (**Figure 6G**). SARS-CoV-2 AIM assays also displayed lower inter-site variability, with CVs significantly <30% for all CD4^+^ and CD8^+^ AIM pairs (**Figure 6H**). ICC analysis demonstrated greater biological than technical variability between donors for all CMV AIMs. SARS-CoV-2-specific AIM assays yielded lower ICCs, particularly for CD8^+^ T cells (**Figure 6I**), notably in line with low variability between donors.

We recently reported that the reproducibility of data from thawed PBMCs is increased by setting viability cut-offs^52^. We thus determined viability at the time of antigen stimulation within inter-site testing samples. Of 72 samples, 43 (60%) were <80%, 19 (26%) were <70% and 8 (11%) were <60% viable at the time of stimulation **(Figure S14A**). There was a strong positive correlation between viability and SI for CD134/CD25 (CD4^+^ T cells) and CD137/CD69 AIM responses (CD4^+^ and CD8^+^ T cells) (**Figures S14B and S14C**). Accordingly, ICC values for CD69/CD137 (but not CD134/CD25) responses could be increased by excluding low-viability samples **(Figures S14D and S14E**). Thus, cell viability may have a substantial influence on AIM responses and should be considered an important experimental factor in AIM assays with cryopreserved PBMCs.

### An automated AIM gating pipeline performs similarly to experienced human analysts

Manual gating is inherently arbitrary and susceptible to bias^83–85^. Differences in AIM gating strategies between laboratories and individual operators may contribute substantially to variability in results^83,84^, particularly among low-frequency or poorly resolved cell populations^85,86^, including antigen-specific T cells. To determine how analyst variability affects AIM data, multi-centre AIM assay data were analyzed by various operators following the same gating strategy. A central analyst (C1) served as the reference, with the other analysts aiming to replicate data from C1. The second central analyst (C2) analyzed data from all sites, while four operators (O1-O4) each analyzed data from one site. Although CVs were generally below 30%, we observed significant and site-dependent variability in SIs between data analysts for both CMV and SARS-CoV-2 AIM assays (**Figures 7A and S15A-D**).

**Figure 7.**
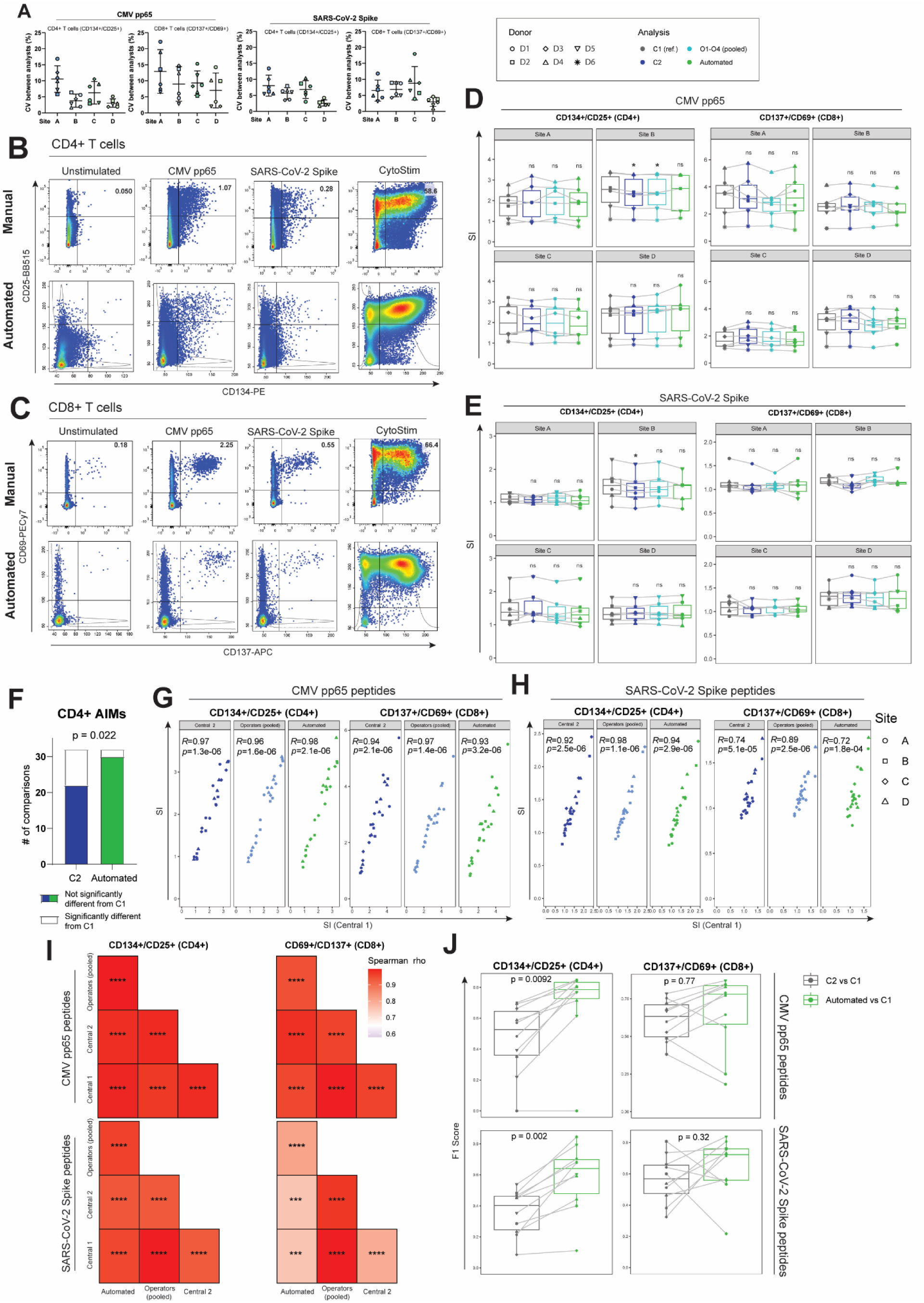
An automated gating pipeline performs comparably to manual analysis for AIM assays. **(A)** Individuals were trained on identical gating protocols and analyzed the same data files from three technical replicates of 20-h AIM assays with CMV pp65 peptides or SARS-CoV-2 Spike peptides on healthy donor PBMCs (n = 6) collected at four distinct sites designated A-D. Raw flow cytometric data from each replicate at each site was analyzed independently by 3 different individuals. Data represent CVs between individual analysts, as a mean of the analyst CV for the three technical replicates of each donor-site combination. **(B and C)** An automated AIM gating pipeline was created via analysis of AIM assay data from healthy donors (n = 6) assayed in triplicate at each of four research centres. Comparative flow cytometric gating approaches for manual and automated analyses are shown for (B) CD4^+^ CD134^+^/CD25^+^ and (C) CD8^+^ CD137^+^/CD69^+^ T cells following a 20-h incubation of cryopreserved PBMCs with no antigen (unstimulated), CMV pp65 peptides, SARS-CoV-2 Spike peptides or CytoStim. The CytoStim condition was used by the automated software to set donor-specific AIM gates, which were then applied to other stimulation conditions for that donor. **(D-F)** Multi-centre CMV and SARS-CoV-2 AIM assay data were analyzed manually or using the automated gating software. A central analyst (C1) defined the gating strategy, provided instructions to the manual analysts (C2 and O1-O4), oversaw automated gating development, and was the reference for comparisons. C1, C2 and the automated software each analyzed all data from all sites, while O1-O4 each analyzed data from a single site (O1: A, O2: B, O3: C, O4: D). (D-F) Box-Cox-corrected AIM SI values are shown for (D) CMV and (E) SARS-CoV-2 Spike AIM assays. P-values were calculated via post-hoc Dunnett’s multiple comparisons test following mixed effects analysis and represent paired comparisons between each analyst and C1, and used to compute (F) the total number of comparisons for C2 and automated analyses that were significantly different from the analysis by C1 for CD4^+^ AIMs. Each of the 32 distinct combinations of CD4^+^ AIM, site and antigen (CMV or SARS-CoV-2) was considered to be one comparison. Pooled analysis of all comparisons was performed using Fisher’s exact test. **(G-I)** Spearman correlations of AIM SI values for C1, C2, O1-O4 (pooled) and automated analyses for CMV and for SARS-CoV-2 Spike. Comparisons are shown against (G and H) the reference C1 and (I) for each analyst against all others in a correlation matrix. (J) F1 scores calculated for C2 manual *vs.* C1 reference (grey) or automated *vs.* C1 reference (green) analysis of CMV and SARS-CoV-2 AIM assay data. Each point represents the F1 score from a unique donor-site combination, with p-values calculated via paired Wilcoxon signed-rank test after averaging F1 scores from technical replicates. ns, not significant (p > 0.05); *, p ≤ 0.05; **, p ≤ 0.01; ***, p ≤ 0.001; ****, p ≤ 0.0001. See also Figures S15-S19.

As automated gating pipelines eliminate operator variability^83,85–87^, we created an automated AIM gating method using our previously described flowDensity approach^88^, which uses nested sequential bivariate gating based on marker density distributions, and applied this to the multi-centre dataset (**Figures 7B, 7C, S16A and S16B)**. Results from the manual and automated analyses were compared to C1. The automated analysis produced SI values that were not significantly different from C1 for all CMV AIM CD4^+^ data and all but two AIM-site combinations for SARS-CoV-2 (for which results from one or more manual analysts also differed from C1) (**Figures 7D, 7E and S17A-C**). When all AIM-site combinations were considered for CD4^+^ T cells, the automated software produced AIM SI values with fewer significant differences from the C1 reference, outperforming the manual analyst C2 (**Figure 7F**).

Notably, AIM SI values from the automated and all manual analysts showed no significant differences from C1 for all CD8^+^ AIM-site combinations for CMV and SARS-CoV-2, suggesting that automation could provide greater benefit in the analysis of CD4^+^ AIMs. Similar to results from manual analysts, automated analysis SIs correlated significantly with those from the reference analyst C1 and other manual analysts (**Figures 7G-I and S18A-E**). To assess the ability of the automated gating pipeline to define individual events as AIM^+^, F1 scores were calculated relative to C1, using F1 scores for C2 vs C1 to enable comparisons across sites. Compared to C2, the automated gating pipeline produced trending or significantly higher F1 scores for defining total CD4^+^ and CD8^+^ T cell populations and AIM^+^ T cells for all CD4^+^ and CD8^+^ AIM pairs (**Figures 7J and S19A-C**). Together, these results highlight the utility of using an automated gating platform for high-throughput analysis of AIM assay data.

## DISCUSSION

Here we developed an optimized AIM assay workflow by systematically testing and addressing key points of technical variation. We demonstrate its reproducibility for the quantification of CMV-specific and SARS-CoV-2-specific T cell responses within and between four research centers. Furthermore, we present the first rigorous analysis of antigen-specific Treg detection in AIM assays and comparison of Treg AIMs in response to diverse clinically relevant antigens. We also describe key advances in the AIM data analysis workflow. First, we developed a Box-Cox transformation-based method to correct for background unstimulated AIM^+^ frequencies, reducing technical variability and enhancing signal detection. Second, we created an automated gating pipeline that eliminates operator variability during AIM flow cytometric data analysis and performs similarly to or better than experienced human analysts. This reproducible workflow will facilitate AIM assay implementation as an efficient and sensitive method for measuring T cell responses in clinical samples.

Reported AIM assay protocols are highly heterogeneous with respect to culture medium, stimulation time, marker selection and data analysis strategies. Illustrating this heterogeneity, the reported vaccine-stimulated frequencies of SARS-CoV-2-specific T cells differed by orders of magnitude between cohorts in a recent meta-analysis of 48 datasets from 12 studies^89^. Using CMV pp65 as a model antigen with strong clinical relevance, we quantified the sources of variability in AIM assays, finding that even when testing technical replicates at the same site, CVs of raw AIM responses were often >50% for CD4^+^ T cell AIMs, consistent with previously reported challenges in reliably detecting rare populations in flow cytometry-based analyses^86,90^. Variation between operators was a substantial contributing factor in the present study, highlighting the need for solutions that reduce technical variability.

In addition to technical variation, methods to normalize stimulated responses to background unstimulated AIM^+^ frequencies differ widely in the literature, with subtraction of, or division by, background AIM^+^ frequencies commonly reported. We found that the relationship between unstimulated and stimulated AIM^+^ frequencies is complex and dataset-dependent, with strong positive associations in some cases and no discernible relationship in others. Thus, arbitrary decisions to subtract or divide by background AIM^+^ frequencies are mathematically inappropriate. We thus developed a universally applicable method to account for unstimulated AIM^+^ T cell frequencies using Box-Cox transformation prior to subtraction. This method approximates linear subtraction or division depending on the user-specified parameter λ. We found that the transformation equation and value of λ can be flexibly adjusted depending on the mathematical properties of individual AIM datasets (**Document S1)**. Researchers can thus tailor the value of λ based on preliminary analyses of the relationship between stimulated and unstimulated AIM^+^ frequencies in the data of interest. Another option is to adopt a λ-value of 0.5, as we have done here, representing a mathematical compromise between subtraction of and division by the unstimulated condition. Our study demonstrates that use of the Box-Cox correction reduces the influence of background AIM^+^ frequencies on the net AIM signal, decreases technical variability and increases signal detection, providing a robust approach to enhance AIM assay reproducibility at the data analysis stage.

Published AIM assay methods report using fresh whole blood or cryopreserved PBMCs. Using whole blood AIM assays as a reference, we demonstrate that CD4^+^ and CD8^+^ T cell AIM responses can be detected in fresh and/or cryopreserved PBMCs but that the accuracy of the data depends on the antigen and markers of interest. Notably, a 24-h delay in processing blood into PBMCs substantially reduces AIM response magnitude. CD8^+^ T cell AIM responses were more strongly affected by suboptimal cell sources, with signal detection in CD107a-containing assays impaired by a 24-h delay and/or cryopreservation particularly for Infanrix. We also note that stimulation index magnitudes are higher in fresh PBMCs than in whole blood, so this could be a preferred approach for rare cell detection. Although fresh blood or PBMCs is clearly the preferred approach, multi-centre cohort studies often require use of cryopreserved samples. For CMV-specific AIM assays, protein and peptide responses remained detectable in cryopreserved PBMCs for all CD4^+^ and CD8^+^ AIM pairs tested, but AIM signals were diminished or lost with a 24-h PBMC processing delay before cryopreservation. In CD4^+^ T cells, CMV pp65-specific AIM responses were generally more efficiently detected using whole protein antigen to stimulate fresh cells (whole blood or PBMCs), while peptide stimulation was superior in cryopreserved PBMCs. Cryopreservation negatively affects APCs^86,91^, which are required for CD4^+^ T cell activation via MHC II, so these preferential CD4^+^ T cell responses to peptides in cryopreserved samples likely reflects loss of APC viability, antigen-processing capabilities and/or co-stimulatory function.

Optimal stimulation time varies considerably between AIMs and may be further influenced by cell source and antigen structure. We show here that 20 hours of stimulation is an optimal compromising time point which allows detection of a diverse selection of AIMs. Although CD154 and CD107a are detected as early as 6 h in some donors, these AIM responses become more homogeneous at 20 h. Notable exceptions were the CD274/CD69 and 6xAIM assays in PBMCs, which produced more reliable results at 44 h. However, 44-h AIM assays come with the increased risk of bystander activation and proliferation^32,33^, leading to overestimation of the antigen-specific response. Overall, we caution against routine use of a 44-h stimulation when the markers of interest are detectable at 20 h.

Antigen-specific Tregs expand following infection or immunization^68,69^, and their roles in antiviral immunity are complex and poorly understood. Few studies have thoroughly investigated AIM responses in Tregs, and no standardized protocols exist for detecting antigen-specific Tregs. We found that FOXP3 is upregulated in a stimulation-dependent manner in CD4^+^ T cells in AIM assays, while FOXP3^+^HELIOS^+^ Treg frequencies remain stable. Thus dual staining for FOXP3 and HELIOS staining is essential to identify antigen-specific Tregs. Previous reports identified antigen-responsive Tregs as CD154^+^ in 24-h AIM assays^78^; however, as CD154 expression is associated with Treg instability and pro-inflammatory functions^79,92^, we sought to identify other markers for antigen-responsive stable FOXP3^+^/HELIOS^+^ Tregs. Although this subset expressed diverse AIMs upon stimulation, antigen-specific cells were best identified as CD134^+^CD137^+^ at 20 h; there was no further enhancement of signal by adding additional AIMs via Boolean combinations. We also confirmed that CD39 was highly expressed in antigen-responsive Tregs^82^, but found that CD39 could not be used as a substitute for FOXP3 and HELIOS. The combination of CD39, CD134, and CD137 only had a positive predictive value of 61-71% to identify antigen-specific Tregs, and CD39 expression was highly variable between donors. However, if fixation and intracellular staining are undesirable, CD39^+^/CD134^+^/CD137^+^ frequencies could provide a rough estimate of antigen-specific Tregs.

Clinical translation of AIM assays for routine monitoring of antigen-specific T cells requires a standardized protocol with a high degree of reproducibility within and between research centres. Using our optimized protocol we assessed the reproducibility (defined by a CV threshold of <30%)^53^ of CD4^+^ and CD8^+^ AIM stimulation indices within and between four distinct research centres located across Canada. We deliberately selected donors with a broad range of AIM responses to CMV and SARS-CoV-2 to evaluate assay reproducibility in people with strong, weak and negligible responses. Although there was a high degree of AIM assay reproducibility both within and between sites, the marker combinations that remained below the between-site CV threshold of 30% were CD134/CD25, CD137/CD69 and CD134/CD137 for CD4^+^ T cells and CD137/CD69 for CD8^+^ T cells, suggesting that these markers display the high degree of reproducibility required for multi-centre studies.

Finally, we demonstrated that manual data analysis is an important source of variability in AIM assay workflows and developed a fully automated gating pipeline to eliminate this variability. The algorithm standardizes AIM gating through nested bivariate segmentation, producing a mathematically defined, unbiased output that more consistently aligned with a reference analyst than did human manual analysts. Furthermore, we demonstrated the ability to use automated gating for AIM data generated with different flow cytometers located at different centres. Although automated analysis platforms have been reported for general flow cytometric characterization of human PBMCs^86–88^, to our knowledge, this is the first automated flow cytometric gating pipeline developed for AIM data analysis and represents a significant advance towards standardizing and streamlining AIM assays for preclinical and clinical applications. It should be noted that automated gating pipelines rely on the consistency of the data acquisition methods, and therefore changes in equipment and/or procedures may require updates in the pipeline since they will affect the density distribution of the data.

Overall, our work provides a comprehensive evaluation of the variability associated with AIM assays and proposes an optimized protocol along with several strategies to mitigate variability, as summarized in **Table 1**. The Box-Cox correction method to normalize AIM responses to the unstimulated control offers a robust and flexible approach to reduce technical variability and quantify antigen-specific responses in a manner that accounts for unpredictable mathematical relationships between background AIM^+^ frequencies and antigen responsiveness. Our data show that for CD4^+^ T cells, AIM assays using CD134/CD25, CD134/CD69, CD137/CD69 and CD134/CD137 can all offer sensitive and reliable detection of antigen-specific cells across a range of conditions, while co-expression of CD137/CD69 optimally detects CD8^+^ T cells and antigen-responsive CD4^+^/FOXP3^+^/HELIOS^+^ Treg responses are best characterized as CD134^+^CD137^+^. These markers are robust across a variety of antigenic stimulants after a 20 h stimulation and yield reproducible results in cryopreserved PBMCs across multiple sites. Nevertheless, other markers may find utility in specific contexts, and future work should continue to apply and validate our methods in AIM assays across a broad range of clinical contexts.

### Limitations of the study

In order to quantify technical variability and test a range of media, cell sources, time courses and experimental conditions, we used relatively few (n = 4-9) donors per experiment to allow a greater number of replicates per donor (up to 12 when considering both intra- and inter-site replicates). However, this is mitigated by the paired and factorial experimental designs used throughout the study. Given the versatility and generalizability of the Box-Cox correction method for AIM stimulation data, there are many possible ways to explore AIM data analysis using this method (see **Document S1)** and it was beyond the scope of the current investigation to attempt all of them; we elected to utilize the simpler Box-Cox-corrected SI (rather than the mSI) with a fixed λ = 0.5. However, the mathematical accuracy of accounting for background AIM^+^ frequencies could conceivably be further improved by selecting a unique λ for each AIM in each experiment, based on the methods presented in **Figure 2** to estimate optimal λ-values. The more robust mSI method, which avoids linear translation at λ = 1, would be highly recommended when selecting higher λ-values (e.g., λ > 0.7; refer to **Document S1** for more details). Another consideration is that the Box-Cox correction method expresses AIM responses as stimulation indices rather than cell frequencies, precluding numerical quantification. If raw AIM^+^ frequencies are required, then Box-Cox transformed stimulation indices could be calculated in parallel to distinguish between unstimulated and stimulated conditions. Finally, we tested AIM methods to detect T cells specific to viral and vaccine antigens; different strategies may be needed to evaluate responses to self antigens in the context of autoimmunity, tumor neoantigens in cancer, or alloantigens in transplantation. In these cases, baseline T cell activation may also be altered. Overall, future work should continue to broadly validate AIM methods for T cells specific to diverse antigens in a context-dependent manner.

## ACKNOWLEDGEMENTS

The authors wish to acknowledge Dr. Lisa Xu of the BC Children’s Hospital Flow Cytometry Core, for her invaluable assistance with flow cytometry data acquisition.

## AUTHOR CONTRIBUTIONS

M.K.L., S.I., J.D.R., J.B., S.L., and S.V. designed the study. J.Q.H., R.G., L.C., L.L. and J.E.F-D. performed experiments. T.H., S.I., J.Q.H., R.G., L.S., L.L. and J.E.F-D. analyzed raw flow cytometry data. G.B. developed the Box-Cox transformation method and wrote Document S1. J.H., T.H., G.B., S.I. and M.K.L. developed and applied the Box-Cox Application to further analyze data. T.H., G.B. and S.I. performed formal analysis of flow cytometry metadata and interpreted results. D.Y., R.B., S.I., T.H. and M.K.L. developed the automated AIM gating software. T.H. and G.B. performed all statistical analyses. T.H. wrote the initial draft of the manuscript. All authors revised and approved the final manuscript for submission. M.K.L. supervised the project and acquired funding.

## STAR METHODS

### KEY RESOURCES TABLE

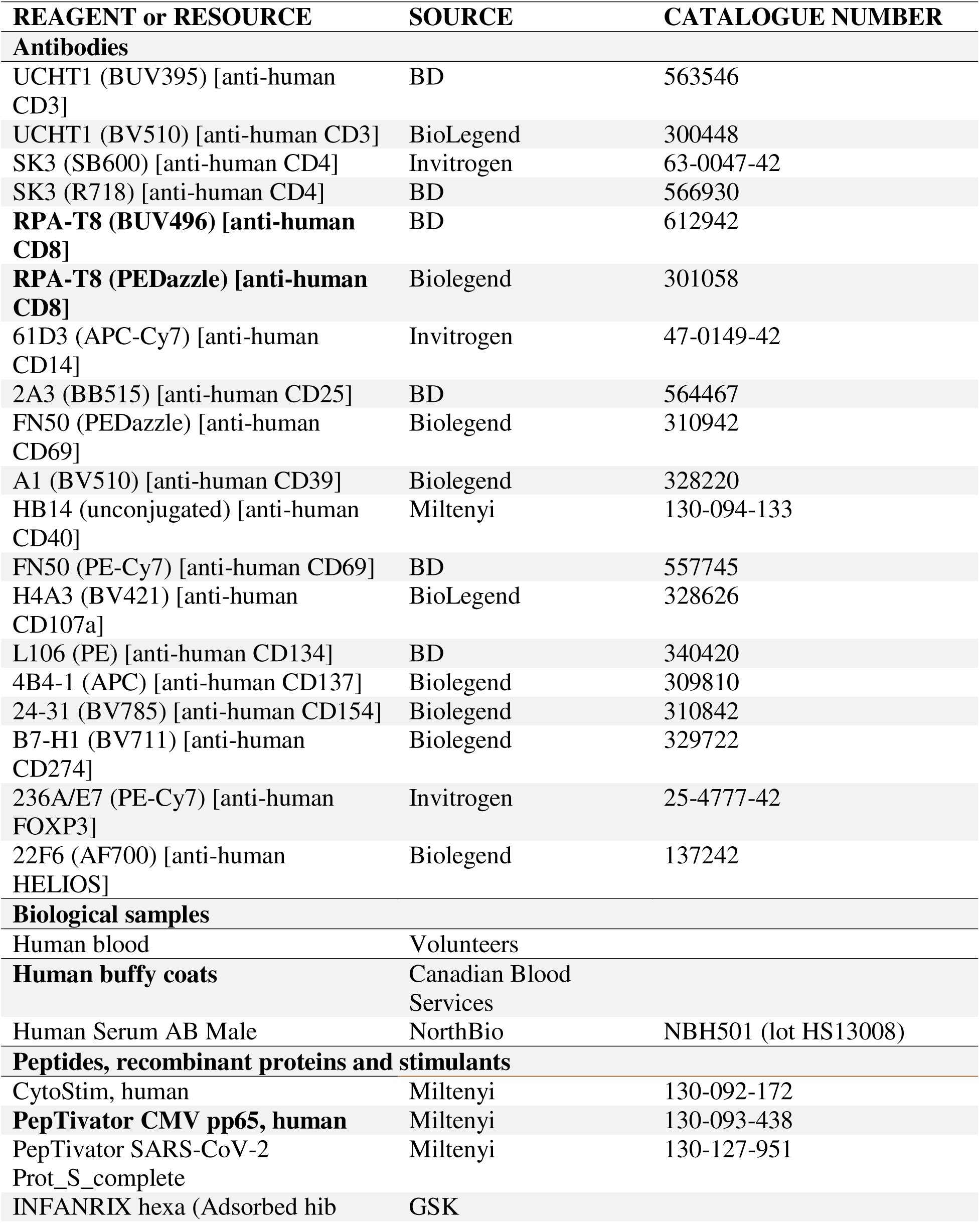
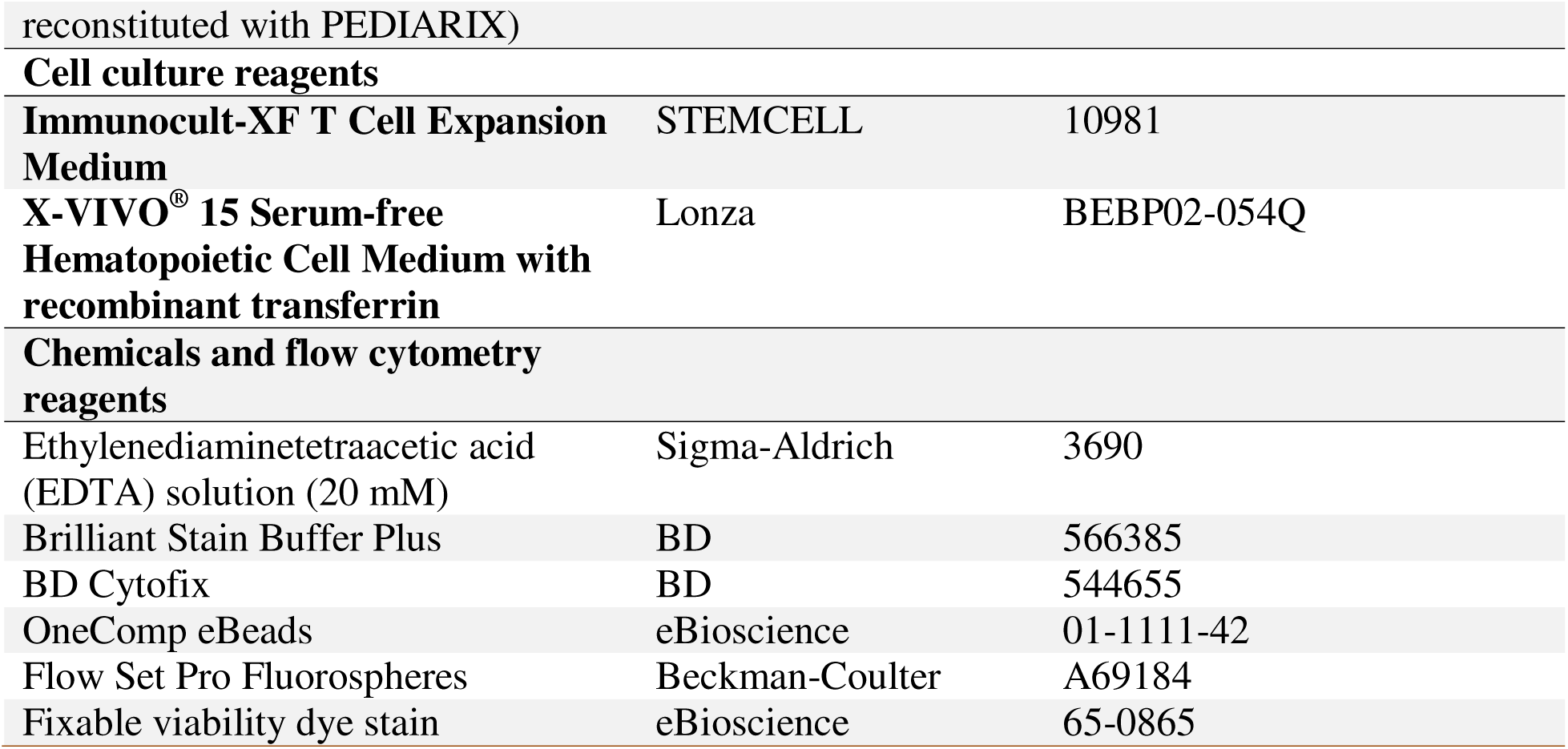

## RESOURCE AVAILABILITY

### Lead contact

All inquiries regarding this work, including requests for materials, reagents and resources, should be directed to and will be addressed by the lead contact, Dr. Megan K. Levings

### Materials availability

Access to biological samples (blood and PBMCs) is limited by the ethics approval for this study. This work did not lead to the generation of new unique products or reagents.

### Data and code availability

Data will be made available upon reasonable request. Code for the Box-Cox correction method and estimation of optimal lambda, along with their application to example AIM datasets, is available on GitHub.

## EXPERIMENTAL MODEL AND SUBJECT DETAILS

### Ethics statement

All work described in this study was performed in agreement with the Declaration of Helsinki and was approved by an institutional review board. Blood samples were provided either as day old buffy coats by Canadian Blood Services CBS REB #2020.038 or from volunteer donors who gave informed consent to participate in the study, UBC Clinical Research Ethics Board # H20-01541.

### Participants and samples

Study participant characteristics are summarized in **Table S1**. **METHOD DETAILS**

### Activation-induced marker (AIM) assay

Peripheral blood samples were collected into sodium heparin vacutainer tubes and either plated immediately for AIM assays or stored at room temperature for 24 h before plating. PBMCs were isolated either from fresh or 24 h blood using density gradient centrifugation and either plated for AIM assays or cryopreserved in CryoStor CS-10. PBMC isolation, cryopreservation and thawing are described in the CAN-ASC consensus protocol on protocols.io (dx.doi.org/10.17504/protocols.io.brrsm56e). Cryopreserved PBMCs for AIM assays were thawed directly into 37°C medium used for stimulation, either Immunocult^TM^-XF (StemCell Technologies), X-Vivo 15 (Lonza), or X-Vivo with 5% human AB serum. For PBMC-based AIM assays, PBMCs were assayed immediately after isolation or, if thawed, after overnight rest at 37°C. 10^6^ live PBMCs were plated in 200 uL medium per well of a round-bottom 96 well plate. For the inter-site testing, viability was determined according to permeability (PI or trypan blue) staining after overnight rest of thawed PBMCs prior to stimulation. PBMCs or 100 µL blood were stained for 15 minutes at 37°C with 1 µg/mL αCD40 blocking antibody, 1/100 CD4-SB600 or CD4-R718 and 1/50 CD107a-BV421 in a final volume of 200 µL prior to stimulation. Cells were incubated for 6, 20 or 44 h with media, 1.5 µg/mL PepTivator CMV pp65 (Miltenyi), 3.3 µg/mL CMV pp65 recombinant protein (Miltenyi), 1 µg/mL PepTivator SARS-CoV-2 Prot_S (Miltenyi), 1/100 Infanrix hexa (GSK) or 1/400 CytoStim (Miltenyi). Prior to harvest, cells were incubated at room temperature for 15 minutes with 2 mM EDTA to detach adherent cells. Cells were stained extracellularly and intracellularly with the appropriate panel of fluorescent-conjugated antibodies for CD3, CD4, CD8, fixable viability dye and AIMs (Tables S1 and S2). Cells were fixed in Cytofix (BD) for 30 minutes at 4°C in the dark, washed and resuspended in PBS for acquisition. Data were acquired on LSR-Fortessa or FACSymphony flow cytometers with Diva v7-9 software (BD) or a CytoFLEX flow cytometer with CytExpert software (Beckman Coulter).

### Data visualization and software development

Graphs were generated with GraphPad PRISM (versions 10.0.0-10.2.3). Figures 1A, 2C and 6A were generated using Biorender.com. The automated gating pipeline was developed using a semi-supervised approach with flowDensity^88^, which identifies the populations based on features detected in the 1D density distribution of the analyzed marker. For this analysis, the transformation functions for the log channels were the same as used in the manual analysis, to maintain consistency in the comparison. The script was created emulating the gating process used in the manual gating by the central analyst C1, using the CytoStim-stimulated condition from each donor to define gates for equivalent samples from the same donor. As the sites used different flow cytometers (two sites used CytoFLEX (Beckman Coulter) and two used LSRFortessa (BD Biosciences)), we created separate pipelines for these two cytometer types that produced similar output data. The Box-Cox correction application was developed using R v4.2.0^93^ with the shiny package^94^ and custom scripts.

## QUANTIFICATION AND STATISTICAL ANALYSIS

### Flow cytometry data analysis

Raw flow cytometry data were analyzed using FlowJo v10.6-10.10 to determine AIM^+^ frequencies among CD4^+^ and CD8^+^ T cells and CD4^+^/FOXP3^+^/HELIOS^+^ Tregs. AIM gates were defined based on the CytoStim-stimulated positive control for each sample and applied across all conditions from that sample. AIM^+^ cells were defined as co-expressing both markers within a pair of AIMs, with Boolean 6xAIM combinations calculated as previously described^49^, such that 6xAIM^+^ cells co-expressed at least one of six AIM pairs: CD134/CD69, CD154/CD69, CD137/CD69, CD134/CD137, CD134/CD154 or CD137/CD154. Boolean quantification of 3xAIM^+^ Tregs was calculated similarly as Tregs expressing at least one of three AIM pairs: CD134/CD69, CD137/CD69, CD134/CD137.

### Statistical methods and transformations

Raw AIM^+^ frequencies were transformed using the Box-Cox formula (see text) with λ = 0.5 (unless otherwise specified), using R (v4.2.0) with custom scripts. Simple statistical analyses and descriptive statistics were performed with GraphPad PRISM (v10.2.2) and R (v4.2.0). CVs were calculated from means and sample standard deviations. When comparing CVs from data with different distributions (e.g., following subtraction of *vs* division by *vs* Box-Cox-correction for the unstimulated condition), an adjusted CV metric was calculated via a bootstrapping approach, where the same transformation was repeatedly applied after randomly replacing unstimulated values. Variance components were computed using a mixed effects model (REML) with donor, operator and replicate as random effects, using R (v4.2.0) with the lme4 package^95^ and custom scripts. Two methods were used to estimate optimal values of λ to minimize correlations between net AIM^+^ values (stimulated – unstimulated, computed after box-cox transformation) and the raw unstimulated AIM^+^ frequency, using three methods (see **Document S1** for additional information):

1. Linear regression, to identify the λ value giving the highest probability of a zero slope (̂β = 0), expressed as a posterior probability distribution for λ
2. Spearman correlation to identify the value of λ resulting in an estimated zero correlation
3. Likelihood profile for λ based on linear regression to estimate the probable optimal value of λ

ICC values (using ICC2) were calculated from variance components obtained by mixed effects analysis (REML) using donor, experiment and site as random effects. Comparisons between vaccination timepoints in solid organ transplant recipients were performed using previously published data^55^, via mixed effects analysis (REML) with Dunnett’s multiple comparisons test.

The effect of media was assessed via mixed effects analysis (REML) with Dunnett’s multiple comparisons test. The effect of cell source on viability was analyzed using the Friedman test with post-hoc planned comparisons via Dunn’s multiple comparisons test. For the effect of cell source on AIM signal detection and for comparison of Treg AIM SIs, 95% CIs were calculated from log_2_-transformed SI data, considering the AIM signal to be significant when the 95% CI did not overlap with zero. The effect of stimulation time on AIM signal detection was assessed by one-sample t-tests on log_2_-transformed SI data, considering the AIM signal to be detectable at each timepoint where the mean log_2_-transformed SI differed significantly from zero. FOXP3^+^ and FOXP3^+^/HELIOS^+^ cell frequencies among CD3^+^/CD4^+^ T cells were compared over time using a two-way repeated measures ANOVA with Tukey’s multiple comparisons test. CD39^+^ cell frequencies between Tconv and Treg subpopulations were compared using a one-way repeated measures ANOVA and Dunnett’s multiple comparisons test. Sensitivity and positive predictive values of CD39 for AIM^+^ Tregs were calculated using frequencies of CD39^+^CD134^+^/CD137^+^/FOXP3^+^/HELIOS^+^/CD3^+^/CD4^+^ cells as the true positive population (CD39^+^ AIM^+^ Tregs), CD39^-^CD134^+^/CD137^+^/FOXP3^+^/HELIOS^+^/CD3^+^/CD4^+^ (CD39^-^ AIM^+^ Tregs) as false negatives, and all other CD39^+^/CD3^+^/CD4^+^ (CD39^+^ Tconvs and CD39^+^ non- AIM^+^ Tregs) and CD39^-^/CD3^+^/CD4^+^ (CD39^-^ Tconvs and CD39^-^ non-AIM^+^ Tregs) T cells as the false positive and true negative populations, respectively:

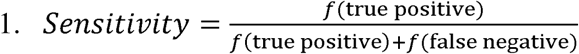

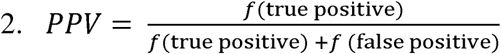

For multi-centre AIM assay data, technical replicates from the same donor at each site were used only to calculate intra-donor CVs, and were averaged prior to statistical analysis. ICCs for multi-centre testing were calculated both between sites after averaging technical replicates, and between technical replicates within sites using the psych package^96^ from R.

A detailed manual gating protocol for analysis of AIM data was developed by the CAN-ASC Central lab (C1) and distributed to operators at all sites; C1 analyses served as the model for development of automated gating software. For comparisons with multiple manual analysts and automated gating, C1 plots were used as the reference. Mixed effects analyses with post-hoc Dunnett’s multiple comparisons were used to compare AIM SI values from manual and automated analysts to the reference, considering each site independently, and these data were used to compare, using Fisher’s exact test, the numbers of total comparisons that were statistically different between (a) the automated software *versus* C1 and (b) C2 *versus* C1. For correlation analyses, data from all sites were pooled and Spearman correlation coefficients were used to assess the relationship between SI values obtained via automated or manual analysis. Wilcoxon signed-rank tests were used to compare F1 scores calculated relative to the reference (C1) for C2 and automated analyses.

