## Supplementary material for "Reproducible detection of antigen-specific T cells and Tregs via standardized and automated activation-induced marker assay workflows": Document S1

**Document S1. The Box-Cox transformation and its application to AIM assay data.**

**Definition of Box-Cox transformation and its inverse function**

The Box-Cox transformation describes a family of power transformations^1,2^. These can be applied to AIM assay data prior to statistical analyses to improve validity of underlying hypotheses, such as normality. The transformation, for strictly positive values of *x,* takes the form:

$$(1)f_{\lambda}\left( x \right)= \left\{ \begin{aligned} \frac{x^{\lambda}-1}{\lambda}\text{, λ}\text{ }\text{≠}\text{ }\text{0,} \\ \log\left( x \right), \lambda=0. \end{aligned} \right.$$

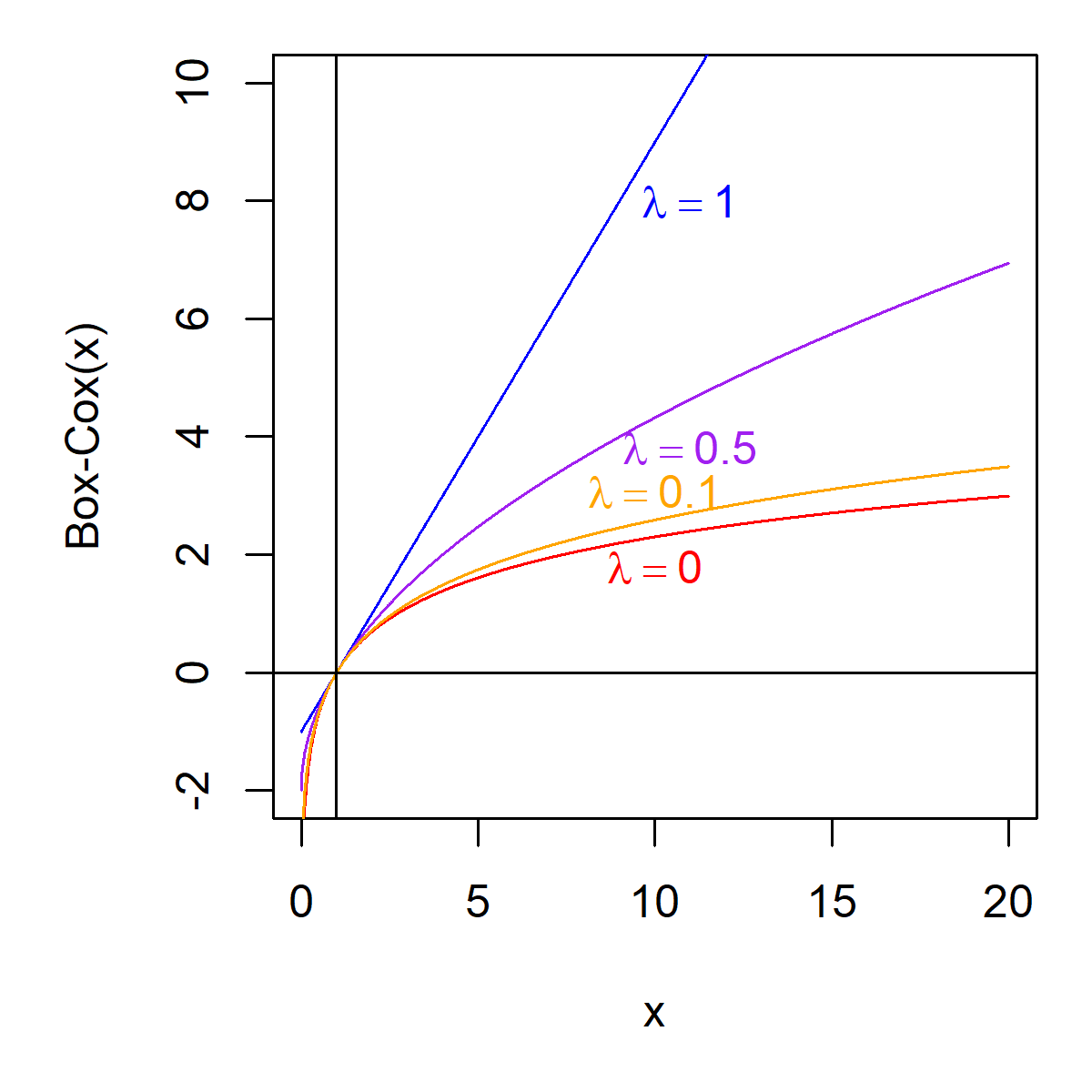

**Figure 1. Illustration of Box-Cox transformation.**

**Figure 1** illustrates the behavior of the Box-Cox transformation for different values of λ. The Box-Cox transformation is equivalent to a simple translation (x-1), when λ=1, and gets more similar (converges) to the logarithm as λ progress to 0.

In mathematical term, it can be shown that:

$$(2)\lim_{\lambda\to0^{+}} \frac{x^{\lambda}-1}{\lambda}=\log\left( x \right).$$

The Box-Cox transformation, with λ close to 0 can then be used as an approximation of the natural logarithm. This approximation is better for values of *x* closer to 1.

The reverse of the Box-Cox transformation is the function that gives us back the original value. This function is noted $f_{\lambda}^{-1}\left( x \right)$ and is defined as:

$$\left( 3 \right) f_{\lambda}^{-1}\left( x \right)= \left\{ \begin{aligned} \left( x\lambda+1 \right)^{\frac{1}{\lambda}}\text{, λ}\text{ }\text{≠}\text{ }\text{0,} \\ e^{x}, \lambda=0. \end{aligned} \right.$$

The reverse Box-Cox is a simple translation (*x*+1) when λ=1 and converges to the exponential function as λ gets closer to 0. It can be shown that:

$\left( 4 \right) \lim_{\lambda\to0^{+}} \left( x\lambda+1 \right)^{\frac{1}{\lambda}}=e^{x}$.

The reverse Box-Cox transformation, with λ close to 0 can then be used as an approximation of the exponential function. This approximation is better for values of *x* closer to 0.

**Box-Cox based operation**

We will define an operation on $x_{1}$ (stimulated) and $x_{0}$ (unstimulated), which is based on the following model:

($SEQ Equation \backslash* ARABIC$ $5$)	$f\left( SI \right)=f\left( x_{1} \right)-f\left( x_{0} \right),$

where $f$ is the Box-Cox transformation ($1$) and *SI* is the measured effect of stimulation. The objective is then to obtain the value *SI* from $x_{1}$ and $x_{0}$.

If we subtract the values $x_{1}$ and $x_{0}$ after Box-Cox transformation, and then use the reverse Box-Cox transformation on the result, we get an operation on values $x_{0}$ and $x_{1}$ defined by the function:

$$\left( 6 \right) g_{\lambda}\left( x_{1},x_{0} \right)=\left\{ \begin{aligned} \left( x_{1}^{\lambda}-x_{0}^{\lambda}+1 \right)^{\frac{1}{\lambda}}, &\text{λ}\text{ }\text{>}\text{ }\text{0} \\ \frac{x_{1}}{x_{0}} , &\text{λ}\text{ }\text{=}\text{ }\text{0}\text{.} \end{aligned} \right.$$

When $\lambda=1$, this operation is equivalent to subtracting the two values, and then adding 1, that is:

$$(7)g_{1}\left( x_{1},x_{0} \right)=x_{1}-x_{0}+1.$$

When $\lambda=0$, this operation is equivalent to the ratio of the two values, as additive operations to log-transformed data are equivalent to multiplicative operations on the original scale.

Let’s consider what is happening when $\lambda$ gets closer to 0.

It can be shown, numerically or via Taylor series expansion, that:

$$(8)\lim_{\lambda\to0^{+}} \left( x_{1}^{\lambda}-x_{0}^{\lambda}+1 \right)^{\frac{1}{\lambda}}=\frac{x_{1}}{x_{0}}.$$

It means that the operation $g_{\lambda}\left( x_{1},x_{0} \right)$ can be used as an approximation of the ratio $\frac{x_{1}}{x_{0}}$, when $\lambda$ is close to 0.

The operation $g_{\lambda}\left( x_{1},x_{0} \right)$ then makes a transition from the ratio $\frac{x_{1}}{x_{0}}$, when $\lambda=0$ to the translated difference $x_{1}-x_{0}+1$, when $\lambda=1$.

There are some limitations to this operation. When $\lambda=0$, $x_{0}$ must be different from zero. Here we will consider strictly positive $x_{1}$ and $x_{0}$. Another limitation of the operation is that it will be well defined, for any $\lambda$, in the interval $x_{1}^{\lambda}-x_{0}^{\lambda}+1\geq0$. Outside of this interval, we might get undefined results, that is, the result can’t be computed.

**Modified Box-Cox based operation**

The translation performed at $\lambda=1$ can be considered a caveat by some users. We can then introduce a correction parameter $\theta$, where $0\leq\theta\leq\text{1}$, and define a more general operation:

$$\left( 9 \right) g_{\lambda,\theta}\left( x_{1},x_{0} \right)=\left\{ \begin{aligned} \left( x_{1}^{\lambda}-x_{0}^{\lambda}+1-\lambda\theta\right)^{\frac{1}{\lambda}}, &\text{λ}\text{ }\text{>}\text{ }\text{0} \\ \frac{x_{1}}{x_{0}} , &\text{λ}\text{ }\text{=}\text{ }\text{0}\text{.} \end{aligned} \right.$$

This operation, when $\theta\neq0$, is no longer the exact application of the Box-Cox transformation and its inverse function. This results in a modification from the initial model ($5$):

$$\left( 10 \right) h\left( SI \right)=f\left( x_{1} \right)-f\left( x_{0} \right),$$

where $f$ is the Box-Cox transformation ($1$), *SI* is the measured effect of stimulation and *h* is a modification to the Box-Cox transformation to:

$$(11)h_{\lambda}\left( x \right)= \left\{ \begin{aligned} \frac{x^{\lambda}-1+\lambda\theta}{\lambda}\text{, }\text{ }\text{ λ}\text{ }\text{≠}\text{ }\text{0,} \\ \log\left( x \right), \lambda=0. \end{aligned} \right.$$

One simple approach to the correction would be to set $\theta=1$. It is to be noted that this approach, for small values of $\lambda,$ is different from the initial formulation, that is, using $\theta=0$, and will converge to the ratio, up to a multiplicative constant:

$$(12)\lim_{\lambda\to0^{+}} \left( x_{1}^{\lambda}-x_{0}^{\lambda}+1-\lambda\right)^{\frac{1}{\lambda}}=\frac{x_{1}}{{e*x}_{0}} .$$

To ensure convergence to the ratio, we suggest the use of a sigmoid function for $\theta\left( \lambda\right)$, such that $\theta\left( 0 \right)=0$ and $\theta\left( 1 \right)=1$. We suggest the following:

$$\left( 13 \right) \theta\left( \lambda\right)=\left\{ \begin{aligned} 1-{(1-\lambda)}^{\left( \frac{\lambda}{1-\lambda} \right)} , &\text{λ}\text{ }\text{<}\text{ }\text{0}\text{.5} \\ \lambda^{\left( \frac{1-\lambda}{\lambda} \right)} , &\text{λ}\text{ }\text{≥}\text{ }\text{0}\text{.5.} \end{aligned} \right.$$

Using this correction makes the $g_{\lambda,\theta}\left( x_{1},x_{0} \right)$ function continuous on $\lambda$ and resolves the translation issue. This more general operation now makes a smooth transition from the ratio

$\frac{x_{1}}{x_{0}}$ ($\lambda\to0^{+}, \theta\to0$) to the difference $x_{1}-x_{0}$ ($\lambda\to1, \theta\to1$).

**Mathematical properties**

For the application to AIM experiments, the operation applied to correct the stimulated value $x_{1}$ by the unstimulated value $x_{0}$ should be decreasing when $x_{0}$ increases. Said otherwise, for a given stimulated value, the operation should return a higher value for lower unstimulated (background) values.

For a given value $x_{1}$and $0<\lambda<1$, $\left( x_{1}^{\lambda}-x_{0}^{\lambda}+1-\lambda\theta\right)^{\frac{1}{\lambda}}$ is indeed decreasing with $x_{0}$ and will reach a minimum (or be undefined) at $x_{0}=\left( {x_{1}}^{\lambda}+1-\lambda\theta\right)^{\frac{1}{\lambda}}$. This can be confirmed by computing the slope i.e. the first derivative:

$$(14) \frac{\partial\left( x_{1}^{\lambda}-x_{0}^{\lambda}+1-\lambda\theta\right)^{\frac{1}{\lambda}}}{\partial x_{0}}=-x_{0}^{\lambda-1}\left( x_{1}^{\lambda}-x_{0}^{\lambda}+1-\lambda\theta\right)^{\frac{1-\lambda}{\lambda}}$$

which has its zero value at $x_{1}^{\lambda}-x_{0}^{\lambda}+1-\lambda\theta=0$ and is negative when $x_{0}<\left( {x_{1}}^{\lambda}+1-\lambda\theta\right)^{\frac{1}{\lambda}}$, for $x_{0}>0$, $x_{1}>0$ and $0<\lambda<1.$

We then need to consider $x_{0}\leq\left( {x_{1}}^{\lambda}+1-\lambda\theta\right)^{\frac{1}{\lambda}}$, which will always be the case when $x_{0}\leq x_{1}$ and $0\leq\theta\leq1$.

**The case of zero and very small values**

The Box-Cox transformation is not defined at $x=0$ when $\lambda=0$, due to the logarithm. For other values of the parameter, $f_{\lambda}\left( 0 \right)=-\frac{1}{\lambda}$.

The operation $g_{\lambda,\theta}\left( x_{1},x_{0} \right)$ is not defined either at $x_{0}=0$ when $\lambda=0$. For small values of $\lambda$, $g_{\lambda,\theta}\left( x_{1},0 \right)$ can be computed:

$$\left( 15 \right) g_{\lambda,\theta}\left( x_{1},0 \right)=\left( x_{1}^{\lambda}+1-\lambda\theta\right)^{1/\lambda}$$

and has its limit to infinity when $x_{0}$ gets closer to 0 ($x_{0}\to0^{+})$. If we consider $x_{1}=1$ and $\theta=0$, we have:

$${(16) g}_{\lambda,\theta=0}\left( 1,0 \right)=2^{1/\lambda},$$

which gives the order of magnitude of the resulting operation, This property allows us to extend the application of the operation $g_{\lambda,\theta}\left( x_{1},x_{0} \right)$ to the case $x_{0}=0$, using $\lambda\neq0$, although $\lambda<0.1$ will results in very large values, as, for example, $2^{1/0.1}=1024$ and $2^{1/0.01}=1.3\times{10}^{30}$.

For small values of $x_{0}<1$, if $x_{0}<x_{1}$, there is a value $\lambda^{*}$ for which $g_{\lambda,\theta}\left( x_{1},x_{0} \right)$ is decreasing with $\lambda\epsilon\left[ 0, \lambda^{*} \right]$. Said otherwise, when using small values for the parameter $\lambda$, the result of the operation gets larger as $\lambda$ decreases. As an example, if $x_{0}=0.01$ and $x_{1}=5$, the ratio would be $g_{\lambda=0, \theta}\left( 5, 0.01 \right)=\frac{5}{0.01}=500$, while $g_{\lambda=0.01, \theta}\left( 5, 0.01 \right)=377.5$ and $g_{\lambda=0.1, \theta}\left( 5, 0.01 \right)=71$.

Using the operation $g_{\lambda,\theta}\left( x_{1},x_{0} \right)$, with a small value of $\lambda\neq0$, then allows the consideration of very low values for $x_{0}$, while reducing the impact of dividing by very small values on the results.

**Application to AIM stimulation data**

We will consider the modified stimulation index (mSI) in its general form to be:

$$(17) {{mSI}_{i}=g}_{\lambda,\theta}\left( s_{i},u_{i} \right)=\left\{ \begin{aligned} \left( s_{i}^{\lambda}-u_{i}^{\lambda}+1-\lambda\theta\right)^{\frac{1}{\lambda}}, &\text{0}\text{ }\text{<}\text{ }\text{λ ≤ }\text{1} \\ \frac{s_{i}}{u_{i}} , &\text{λ}\text{ }\text{=}\text{ }\text{0}\text{,} \end{aligned} \right.$$

where $s_{i}$ is the post stimulation measure $u_{i}$ is the no stimulation measure. We propose to use:

$$(18) \theta\left( \lambda\right)=\left\{ \begin{aligned} 1-(1-{\lambda)}^{\left( \frac{\lambda}{1-\lambda} \right)} , &\text{λ}\text{ }\text{<}\text{ }\text{0}\text{.5} \\ \lambda^{\left( \frac{1-\lambda}{\lambda} \right)} , &\text{λ}\text{ }\text{≥}\text{ }\text{0}\text{.5.} \end{aligned} \right.$$

The modified stimulation index is well defined when: $u_{i}\leq\left( {s_{i}}^{\lambda}+1-\lambda\theta\right)^{\frac{1}{\lambda}}$ and $0\leq\theta\leq1$.

In its simplified form, the stimulation index (SI) has parameter $\theta=0$ and can be expressed:

$$(19) {{SI}_{i}=g}_{\lambda}\left( s_{i},u_{i} \right)=\left\{ \begin{aligned} \left( s_{i}^{\lambda}-u_{i}^{\lambda}+1 \right)^{\frac{1}{\lambda}}, &\text{0}\text{ }\text{<}\text{ }\text{λ ≤ }\text{1} \\ \frac{s_{i}}{u_{i}} , &\text{λ}\text{ }\text{=}\text{ }\text{0}\text{.} \end{aligned} \right.$$

The use of parameter $\theta=0$ is equivalent to the application of Box-Cox transformation and its reverse, as described in STAR Methods.

The use of the $\lambda\theta\left( \lambda\right)$ correction makes the SI more general and is the suggested approach when using larger values of $\lambda$.

To be noted, the stimulation index is proposed for situations when we expect stimulated values to be larger than unstimulated values. When the values are outside the application range, that is, $u_{i}>\left( {s_{i}}^{\lambda}+1-\lambda\theta\right)^{\frac{1}{\lambda}}$, we suggest replacing the result by a small value.

When $\lambda$ is small and $\theta=0$, or as defined above in $(18$), the stimulation index approximates the ratio $\frac{s_{i}}{u_{i}}$, while reducing the impact of very small unstimulated values. Measures from unstimulated conditions in AIM experiments are sometimes small, in a range where the error rate can be important. Using the ratio stimulated/unstimulated can then result in extreme values, not representative of biology. The use of the SI with a relatively small $\lambda$ (e.g. 0.1) can reduce this impact while still accounting for some multiplicative effects between stimulated and unstimulated cells.

A complementary method to deal with very small unstimulated values while computing ratios is to replace the very small values or add a small $\varepsilon$ to all data. This $\varepsilon$ needs to be determined by the analyst, based on the distribution of values, knowledge about the measured parameter and technical considerations, such as the number of cells in parent gate. Combining the stimulation index with the addition of a small $\varepsilon$ would further reduce the impact of very small unstimulated values.

**The issue of scaling**

The result of the stimulation index calculation is dependent on the scale of the data and the choice of parameters λ and θ.

Let’s first consider the generalised version of SI, with an example. Let’s consider a stimulated value of 5% and unstimulated value of 0.1%. We could express these as $s_{i}=5$, $u_{i}=0.1$ or $s_{i}=0.05$, $u_{i}=0.001$. Computing the ratio ($\lambda=0$) will return in both case a value of 50, while the subtraction ($\lambda=1$) would return 4.9 and 0.049, respectively. Ratio is insensitive to scaling, while subtracting is directly proportional to the scale. Between these, the effect of scaling is non-linear. For $\lambda=0.5$, we get stimulation index values of 5.86 and 0.49, respectively

If we now consider the stimulation index with $\theta=0$and $\lambda=1$, the operation produces a translation of the scale of the data ${SI}_{i}=s_{i}-u_{i}+1$. If we suppose a small scale for $s_{i}$ and $u_{i}$, then ${SI}_{i}=s_{i}-u_{i}+1$will be close to 1, for all samples. If we want to compare samples on an additive scale, the difference between samples would not be affected by the translation. If we are investigating fold changes between samples, these would be reduced, compared to using $s_{i}-u_{i}$. Using ${SI}_{i}=s_{i}-u_{i}+1$ would also result in smaller coefficient of variation, due to the translation alone, but also to reduced fold-change variation between individuals. For this reason, we suggest adjusting the θ parameter, and/or scaling the data before computing the stimulation index, when $\lambda$ is closer to 1.

Using the correction proposed in $(18$) solves the translation issue, while ensuring convergence to the ratio at $\lambda\to0^{+}$.

**Choice of** $\boldsymbol{\lambda}$

The choice of $\lambda$ is dependent on the experiment. If we observe or expect a multiplicative relationship between the unstimulated and stimulated data, a choice of $\lambda$ close to 0 is recommended. Use of a small, non-zero value for $\lambda$, such as 0.1, will reduce the impact of very small observations for unstimulated values.

If we observe or expect the unstimulated values to be simple background, a choice of $\lambda$ close to 1 is recommended.

The reality is that different factors can result in different relationships between unstimulated and stimulated values. Batch effect, unexpected activation of cells, difference in reactive, cell survival after freeze/thaw etc. can have unexpected effects on the experiment. When the relationship between unstimulated and stimulated is not clear, a choice of $\lambda=0.5$ is suggested.

**Guidance for the choice of** $\boldsymbol{\lambda}$

We implemented a tool to guide the choice of $\lambda$, based on model ($5$), and the assumption that S should be non correlated with the unstimulated data $x_{0}$. Said otherwise, the objective is that the correction made to the stimulated data $x_{1}$ to obtain the effect of stimulation is effective at removing artifacts or background effects. We used different strategies to identify the optimal $\lambda$, all based on the hypothesis of independent observations.

Method 1: Spearman correlation. Under the assumptions above, the optimal $\lambda$ should give:

$$cor\left( f_{\lambda}\left( S \right)-f_{\lambda}\left( U \right), f_{\lambda}\left( U \right) \right)=0,$$

where $S=(s_{1},s_{2},\ldots,s_{n})$ and $S=(u_{1},u_{2},\ldots,u_{n})$ are the *n* IID (independent, identically distributed) measures for the stimulated and unstimulated conditions, respectively. A positive residual correlation indicates an under-correction from the unstimulated data, while a negative correlation indicates an over-correction. Use of Spearman correlation $\rho$makes the method more robust to outliers than Pearson $r$ and is non-parametric.

Method 2: the beta coefficient

Under a linear regression framework, we can consider:

$f_{\lambda}\left( S \right)= \alpha_{\lambda}+ \beta_{\lambda}{\cdot f}_{\lambda}\left( U \right)+\varepsilon$

Where $\beta=1$ for the optimal $\lambda$. We can then proceed to linear regression, using the Box-Cox transformation with a set of values for $\lambda$ and search for the $\hat{\lambda}$ value such that the estimated slope $\hat{\beta}_{\lambda}=1$.

Method 3: the posterior probability distribution

Under a Bayesian framework for the linear model described above, with a non-informative prior and under normality assumptions, it can be shown that the posterior distribution of $\beta_{\lambda} |S$ follows a t-distribution with estimated mean $\hat{\beta}_{\lambda}$ (the estimated slope from the regression) and scale parameter ${{SE}_{\hat{\beta}}}^{2}$ (the estimated standard error)^3,4^. From this, we can search for an optimal $\hat{\lambda}$ which will maximise the posterior density for $\beta_{\lambda}=1 |S$.

Method 4: the likelihood function

If we suppose the values to be normally distributed, then we can compute the likelihood of $\lambda$, which is the probability of the observed dependent variable *S* under the transformation with parameter $\lambda$ and the linear model above, where we fixed $\beta=1$. This likelihood is obtained from the probability of the transformed data $f_{\lambda}\left( S \right)$, multiplied by the Jacobian of the transformation^5^. For practical reasons, we prefer to work with the maximised log-likelihood, which is, after simplification and up to a constant, and according to Box and Cox^1^ (1964):

$$\mathcal{L}\left( \lambda\right)=-\frac{1}{2}n \log\left( \hat{\sigma}_{\lambda}^{2} \right)+(\lambda-1)\sum_{i=1}^{n} log(s_{i})$$

Where $\hat{\sigma}_{\lambda}^{2}$ is the sum of squared residuals from the model, divided by *n,* and the second term is the log-transformed Jacobian. The optimal $\lambda$ is the value which maximises $\mathcal{L}\left( \lambda\right)$, with 95% confidence intervals approximated by

$$\mathcal{L}\left( \hat{\lambda} \right)\mathcal{-L}\left( \lambda\right)< \frac{1}{2}\chi_{1}^{2}\left( 0.05 \right).$$
