## Supplementary Figures and Tables for "Reproducible detection of antigen-specific T cells and Tregs via standardized and automated activation-induced marker assay workflows"

*Equal contribution

**Co-senior authors

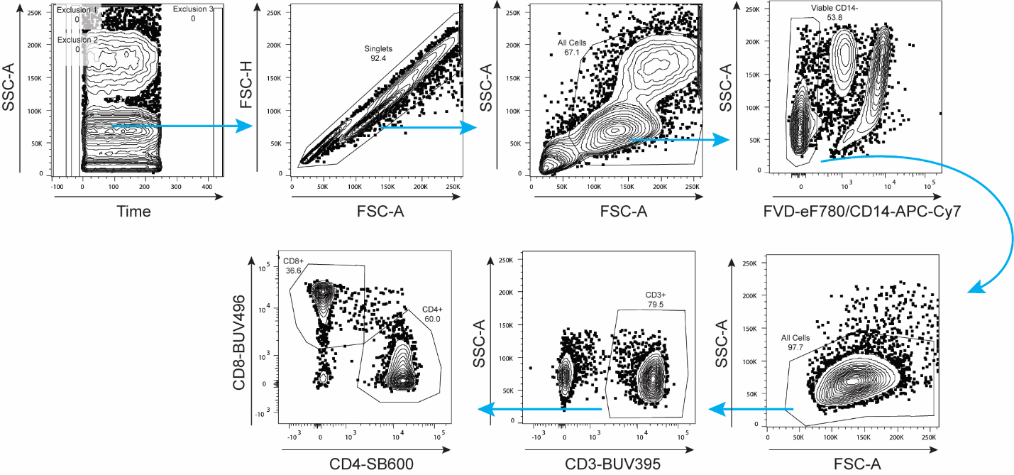

**Figure S1. Representative flow cytometric gating strategy.**

AIM assays were analyzed by flow cytometry to quantify antigen-specific T cell responses. Live CD3^+^ lymphocytes were gated as CD4^+^ or CD8^+^ T cells, with AIM^+^ populations defined downstream of these gates.

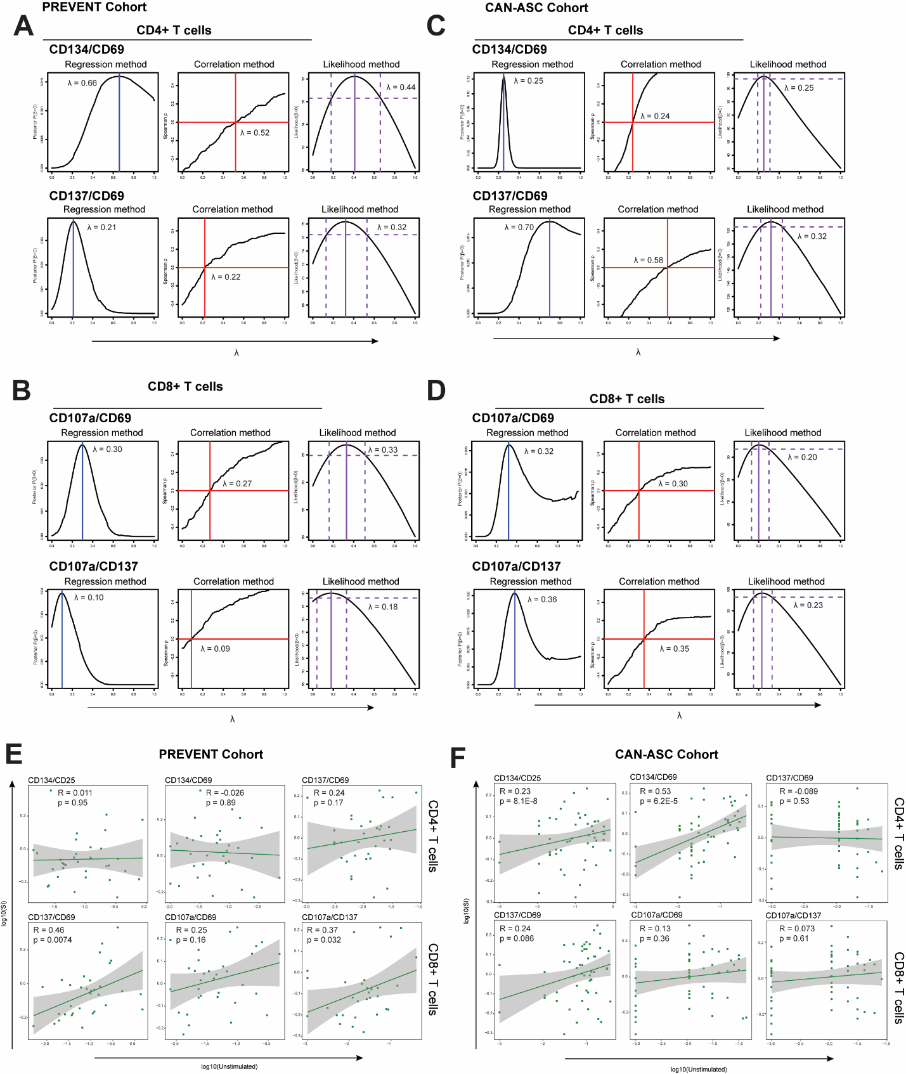

**Figure S2. Optimal values of λ for Box-Cox correction of AIM assay data vary by AIM and between cohorts.**

Analyses of additional AIMs from the PREVENT and CAN-ASC cohorts were performed to validate methods of optimal λ value determination and Box-Cox correction. (**A-D**) Three statistical methods were used to estimate the optimal λ value for the indicated CD4^+^ (A and C) and CD8^+^ (B and D) AIMs, seeking to minimize the correlation between the unstimulated and net stimulated AIM values. Left panel: linear regression, to identify the λ value giving the highest probability of a zero slope ($\hat{\beta}=0$), expressed as a posterior probability distribution for λ. Centre panel: Spearman correlation to identify the value of λ resulting in an estimated zero correlation. Right panel: likelihood profile for λ based on linear regression to estimate the probable optimal value of λ. (**E and F**) Correlation and linear regression analyses of unstimulated AIM^+^ frequencies and Box-Cox-corrected stimulation indices (SI), using λ = 0.5. CD134^+^/CD25^+^ frequencies among CD4^+^ T cells (upper panel) and CD137^+^/CD69^+^ frequencies among CD8^+^ T cells (lower panel) are presented. Spearman ρ and p-values are shown from Spearman correlation tests. The solid line is derived from linear regression analysis. (A-F) Data represent SARS-CoV-2 Spike peptide AIM assays for n = 33 solid organ transplant recipients after three doses of a COVID-19 mRNA vaccine in the PREVENT cohort (A, B and E) or CMV pp65 peptide AIM assays of 6 healthy donors assayed across 3 independent experiments at each of 4 distinct research centres in the CAN-ASC cohort (n = 52 total) (C, D and F). Related to Figure 2.

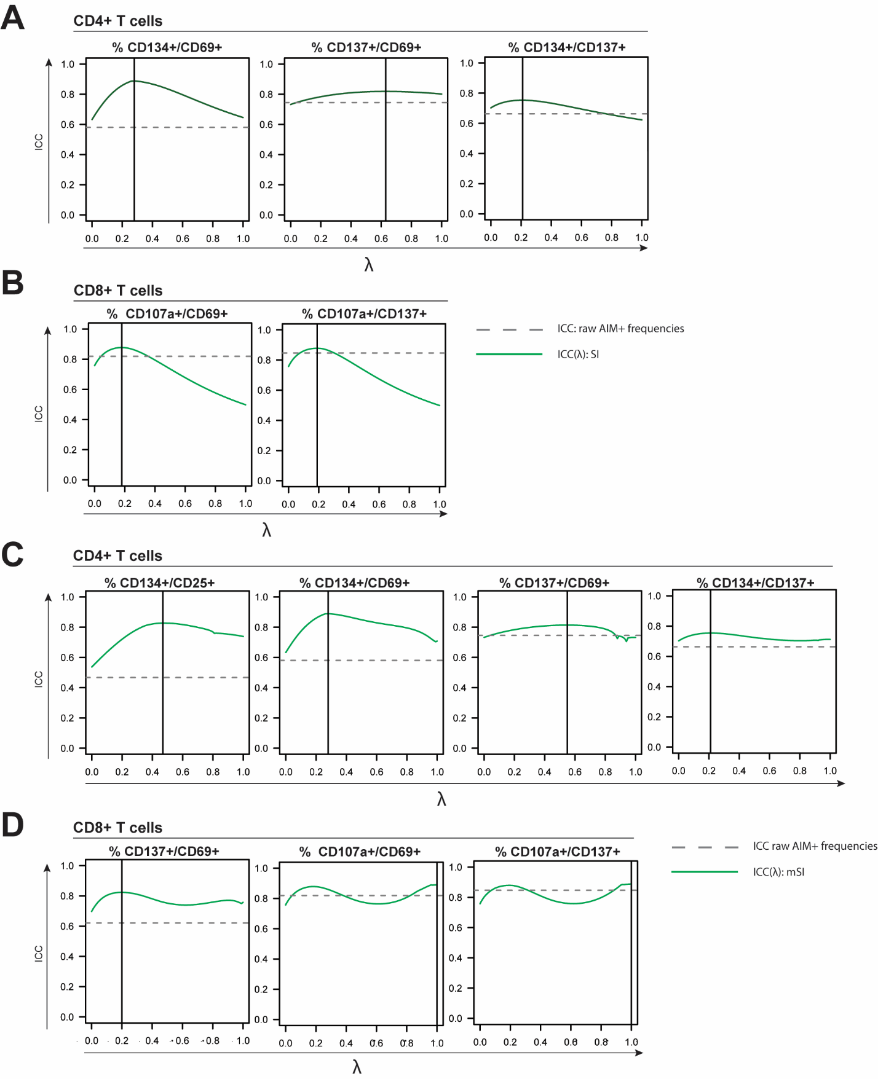

**Figure S3. A Box-Cox-corrected AIM stimulation index enhances detection of biological variability.**

(**A-D**) Ratios of biological (between-donor) variance to total variance (intra-class correlation, ICC; green solid line) plotted as a function of λ using the Box-Cox-corrected SI (A and B) or the scale-independent modified (m)SI (C and D) for CD4^+^ (A and C) and CD8^+^ (C and D) AIM responses to CMV pp65. The maximum value of the function is indicated by a vertical line. When λ approaches 0, the function approximates division by the unstimulated AIM^+^ frequency; when λ approaches 1, the function approximates subtraction of the unstimulated AIM^+^ frequency. ICC2 values for raw (untransformed) AIM^+^ frequencies are indicated by the grey dotted line. Data represent 12 technical replicates collected across four distinct sites for each of n=6 healthy donors. Related to Figure 2. See also Document S1.

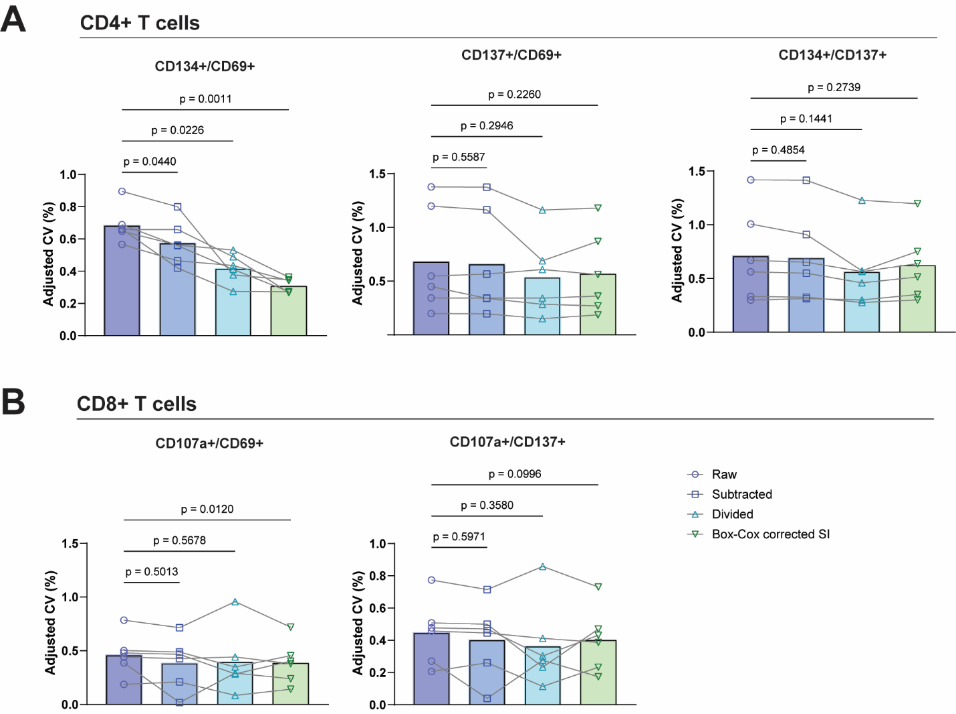

**Figure S4. Box-Cox-corrected SI reduces intra-donor technical variability for some CD4^+^ and CD8^+^ AIMs.**

(**A-B**) Comparison of bootstrapped distribution-robust adjusted CVs calculated between technical replicates within individual donors for raw pp65-stimulated AIM^+^ frequencies among CD4^+^ (A) and CD8^+^ (B) T cells, AIM^+^ responses with subtraction of, or division by, the respective unstimulated AIM^+^ frequency, and Box-Cox-corrected SI. Each point is the CV of technical replicates from one donor, representing within-donor technical variability. P-values are shown from Dunnett’s multiple comparisons test following one-way repeated measures ANOVA. Data are from n = 6 healthy donors assayed in 12 technical replicates. Related to Figure 2.

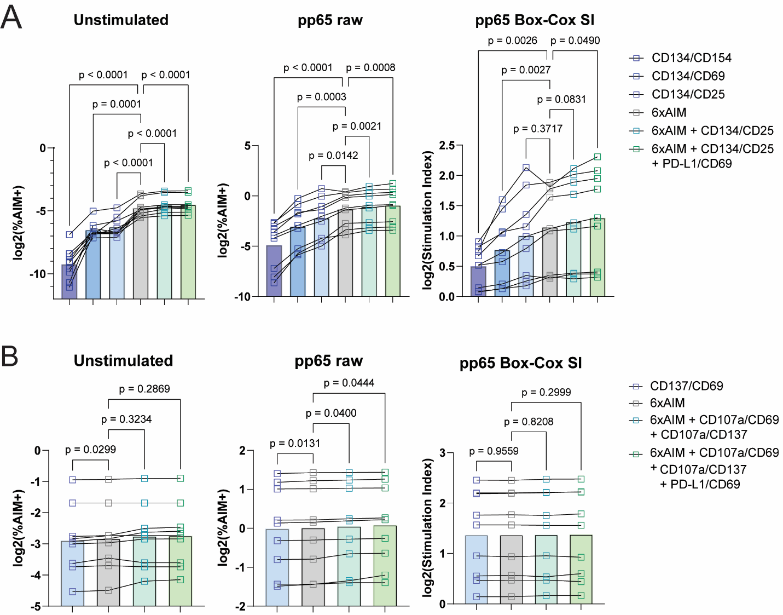

**Figure S5. Box-Cox transformation-based correction for unstimulated AIM frequencies can be used with Boolean analysis to enhance detection of antigen-specific T cells.**
(**A-B**) Antigen-specific T cell detection using Boolean analysis of AIM combinations among CD4^+^ (A) or CD8^+^ (B) T cells in unstimulated or cytomegalovirus (CMV) peptide pp65-stimulated PBMCs following a 20-h incubation. Raw AIM^+^ frequencies in unstimulated (left) and pp65-stimulated (middle) conditions are compared with corrected AIM responses using the Box-Cox transformation-based method (right). Data are from n = 9 healthy donors, with p-values determined by Dunnett’s multiple comparisons test following repeated-measures one-way ANOVA on log_2_-transformed data. Related to Figure 2.

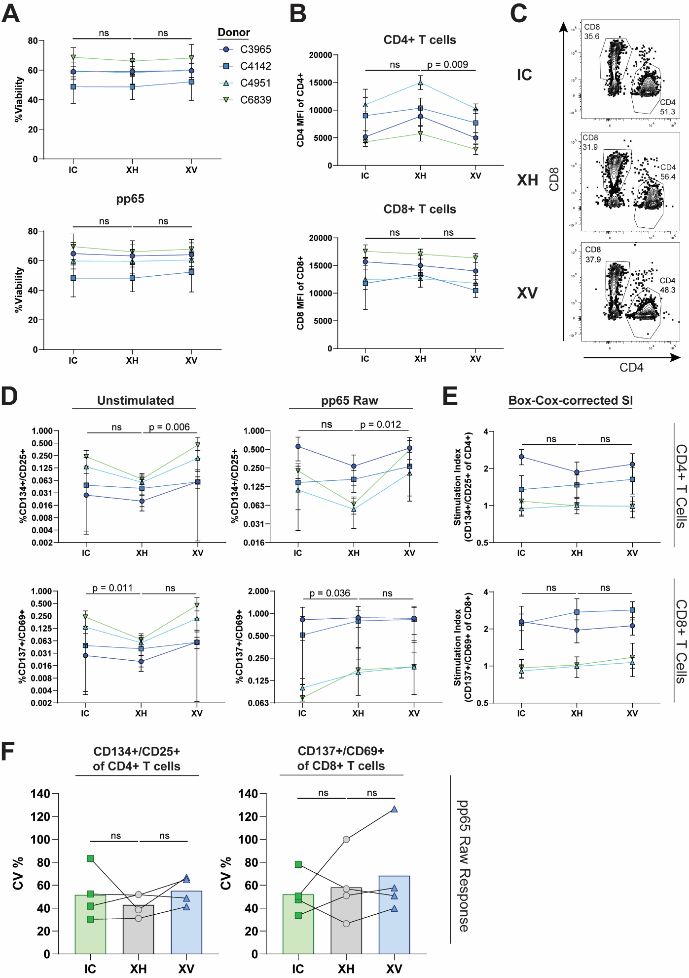

**Figure S6. Serum-free media enables antigen-specific T cell detection without increasing variability.**

**(A)** Percentage of viable CD3^+^ T cells in serum-free ImmunoCult-XF (IC), X-Vivo 15 (XV), or XV + 10% human serum (XH). Cells were left unstimulated (top) or stimulated with the CMV pp65 (bottom). (**B and C**) Geometric mean fluorescence (MFI) of CD4 and CD8 among CD4^+^ and CD8^+^ T cells, respectively (B) and representative flow cytometry plots of CD4^+^ and CD8^+^ frequencies (C) among CD3^+^ T cells in the AIM assay pp65 peptide-stimulated condition. (**D**) Frequencies of CD134^+^/CD25^+^ events among CD4^+^ T cells (top) and CD137^+^/CD69^+^ events among CD8^+^ T cells (bottom) in the AIM assay unstimulated (left) or pp65-stimulated (right) conditions. (**E**) Box-Cox-corrected SI for CD4^+^ (top) and CD8^+^ (bottom) AIM responses following pp65 stimulation for assays conducted in IC, XH and XV. (**F**) Comparison of CVs calculated from raw CD134^+^/CD25^+^ frequencies among CD4^+^ T cells (left) and CD137^+^/CD69^+^ frequencies among CD8^+^ T cells (right) between AIM assays performed in IC, XH and XV. (A, C-E) Shown are p-values from Dunnett’s multiple comparison tests following one-way repeated-measures ANOVA on raw (A, C and F) or log_2_-transformed (D and E) data after averaging technical replicates within each donor: ns, not significant. Error bars represent the standard deviation among technical replicates for each donor. (A-F) Data represent 5-6 technical replicates from n = 4 healthy donors.

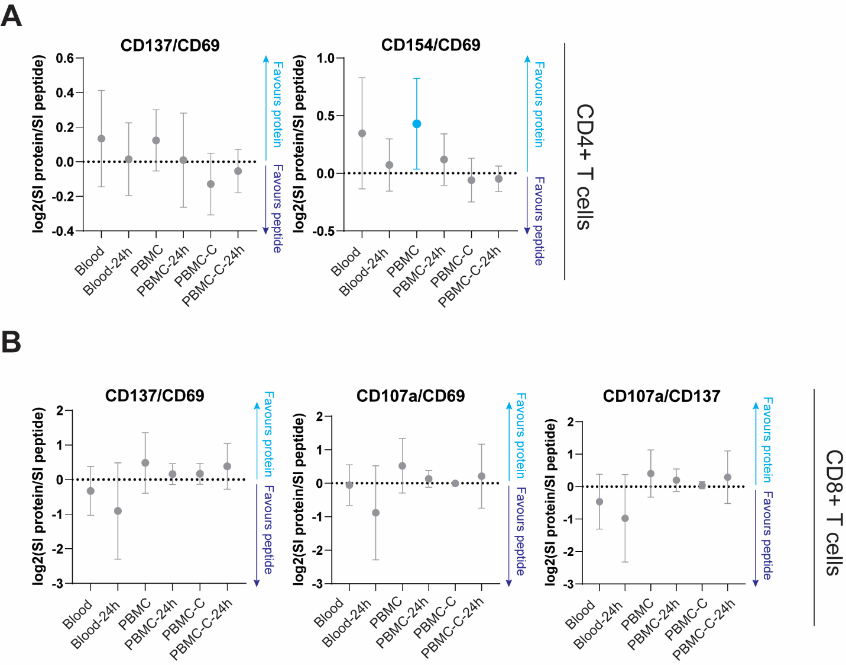

**Figure S7. CD8^+^ AIM detection is similar using whole protein or peptide stimulation.**

Antigen-specific T cell responses following 20-h stimulation with CMV pp65 whole protein or peptide pool, using fresh whole blood, or fresh (PBMC) or cryopreserved (PBMC-C) peripheral blood mononuclear cells, with or without a 24-h delay in processing (-24h). (**A and B**) Ratio of AIM stimulation indices between CMV pp65 protein- and peptide-stimulated CD4^+^ (A) and CD8^+^ (B) T cells in relation to cell source. Positive and negative ratios indicate greater detection of AIM responses with whole protein and peptide stimulation, respectively, while confidence intervals overlapping zero indicate no significant difference between protein and peptide stimulation. Data are from n = 6 healthy donors. Related to Figure 3.

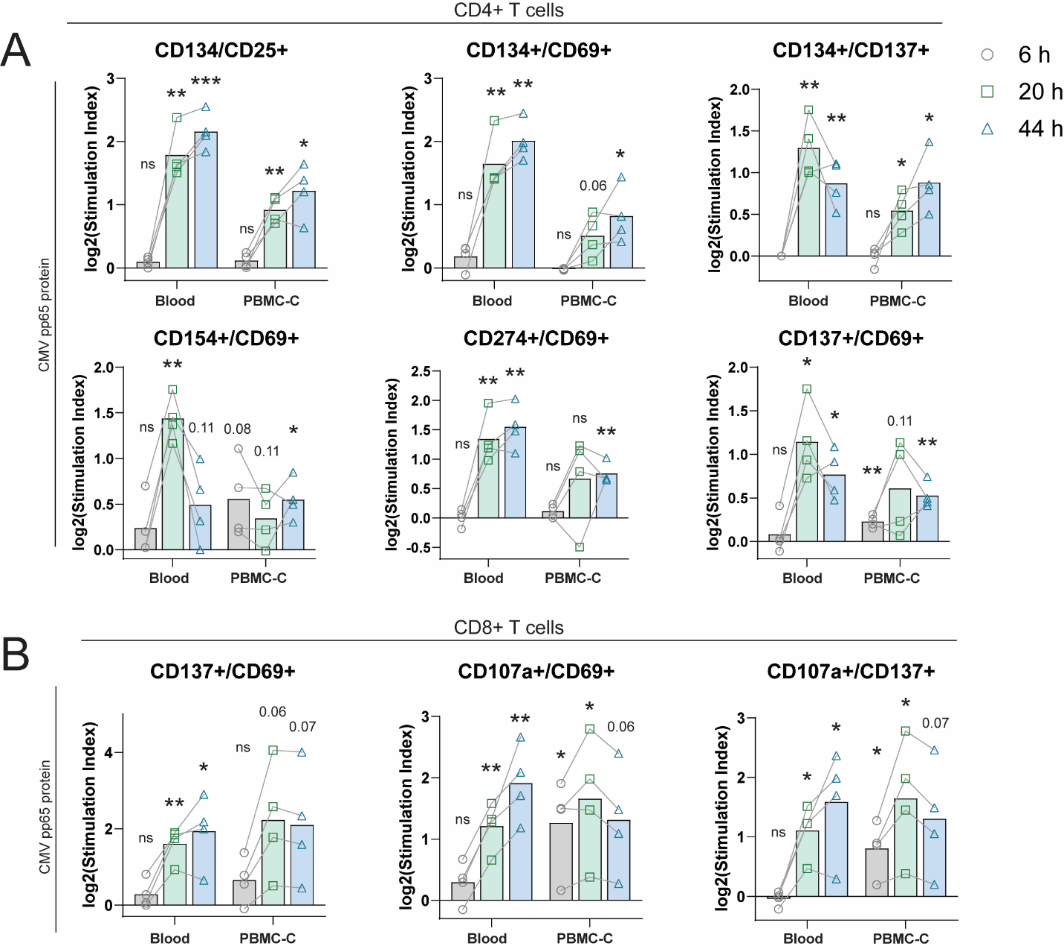

**Figure S8. AIM responses to whole-protein stimulation are optimally detected at 20 h.**

(**A and B**) AIM expression on (A) CD4^+^ T cells and (B) CD8^+^ T cells after stimulation of fresh whole blood or cryopreserved PBMCs (PBMC-C) with CMV pp65 whole protein for 6, 20 or 44 hours. SIs are represented as log_2_-transformed data. P-values are shown from one-sample t-tests, with AIM signals considered present when the mean log_2_-transformed SI significantly differs from zero. All data are paired samples from n = 4 healthy donors. Related to Figure 4.

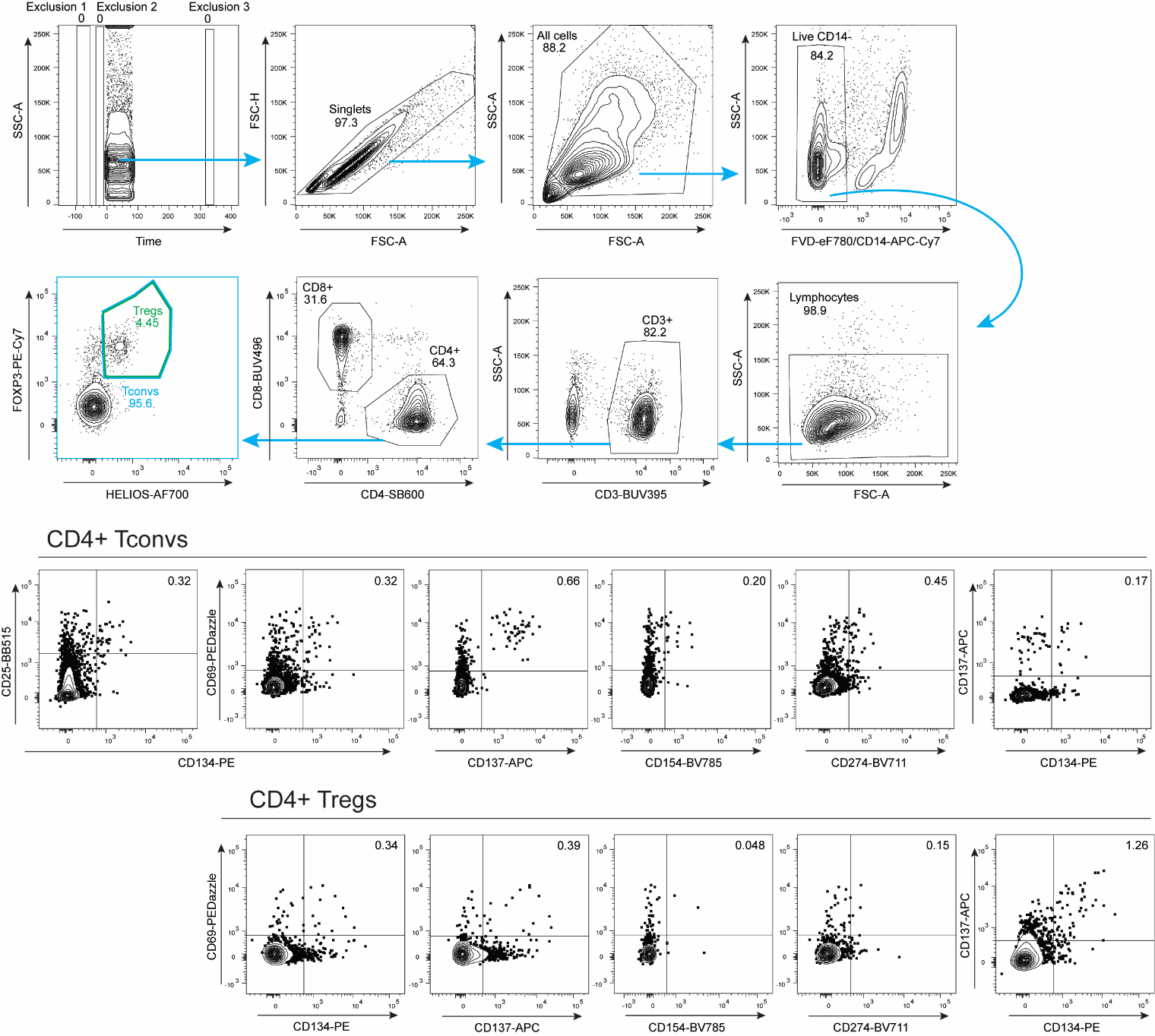

**Figure S9. Representative gating strategy for Tregs in AIM assays.**

AIM assays were analyzed by flow cytometry following to quantify antigen-specific Treg responses. Live CD3^+^ lymphocytes were gated as CD4^+^ or CD8^+^ T cells, and CD4^+^ T cells were further divided into Tregs (FOXP3^+^/HELIOS^+^) and Tconvs (non-Tregs). AIM^+^ Tconvs were defined as CD134^+^/CD25^+^, CD134^+^/CD69^+^, CD137^+^/CD69^+^, CD154^+^/CD69^+^, CD274^+^/CD69^+^ or CD134^+^/CD137^+^. Several AIM pairs were evaluated to define antigen-specific Tregs, including CD134/CD69, CD137/CD69, CD154/CD69, CD274/CD69 and CD134/CD137. Representative data shown are from cryopreserved healthy donor PBMCs stimulated for 20 h with CMV pp65 peptides.

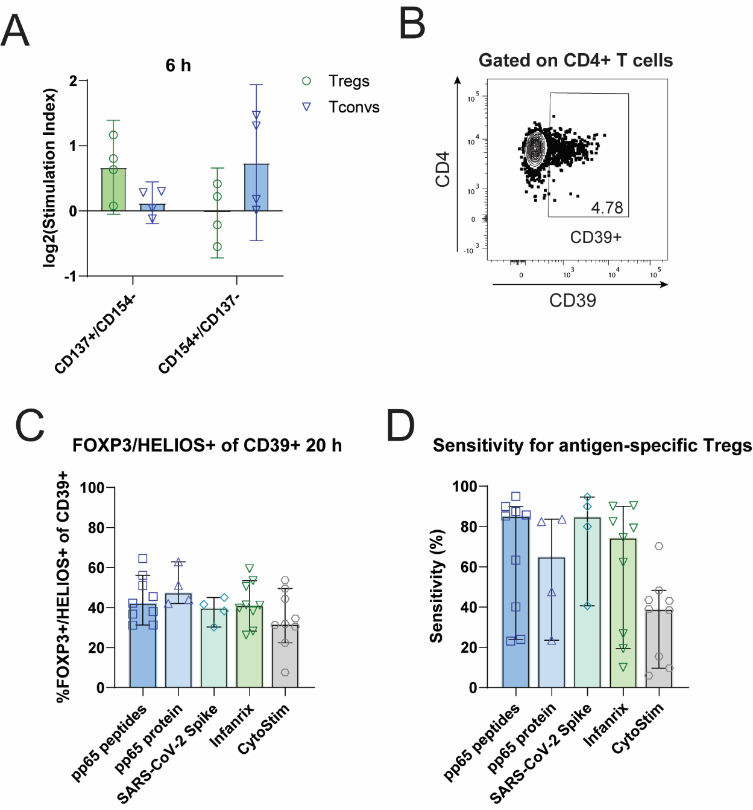

**Figure S10. CD39^+^ and 6-h CD137^+^/CD154^-^ phenotypes variably detect antigen-specific Tregs in AIM assays.**

(**A**) Stimulation index for CD137^+^/CD154^-^ and CD154^+^/CD137^-^ populations among CD4^+^ FOXP3^+^/HELIOS^+^ regulatory T cells (Tregs) and FOXP3/HELIOS^-^ conventional CD4^+^ T cells (Tconvs) after stimulating cryopreserved PBMCs for 6 h with cytomegalovirus (CMV) pp65 peptides. Stimulation index was determined by normalizing pp65 peptide-stimulated AIM^+^ frequencies to the unstimulated control using the Box-Cox correction method described in Figure 2. The mean log_2_-transformed stimulation index is shown for each AIM. Error bars represent 95% confidence intervals of the mean. (**B**) Representative flow cytometric gating on CD39^+^ events among CD4^+^ T cells. (**C**) Quantification of FOXP3^+^/HELIOS^+^ Treg frequencies among CD4^+^/CD39^+^ T cells from PBMCs left unstimulated or stimulated with pp65 peptides for 6, 20 or 44 h.(**D**) Sensitivity of CD39 as a marker for CD134^+^/CD137^+^/FOXP3^+^/HELIOS^+^ Tregs among total CD4^+^ T cells (see methods for details). Data are from n = 4 (A) or n = 9 (B and C) healthy donors. Related to Figure 5.

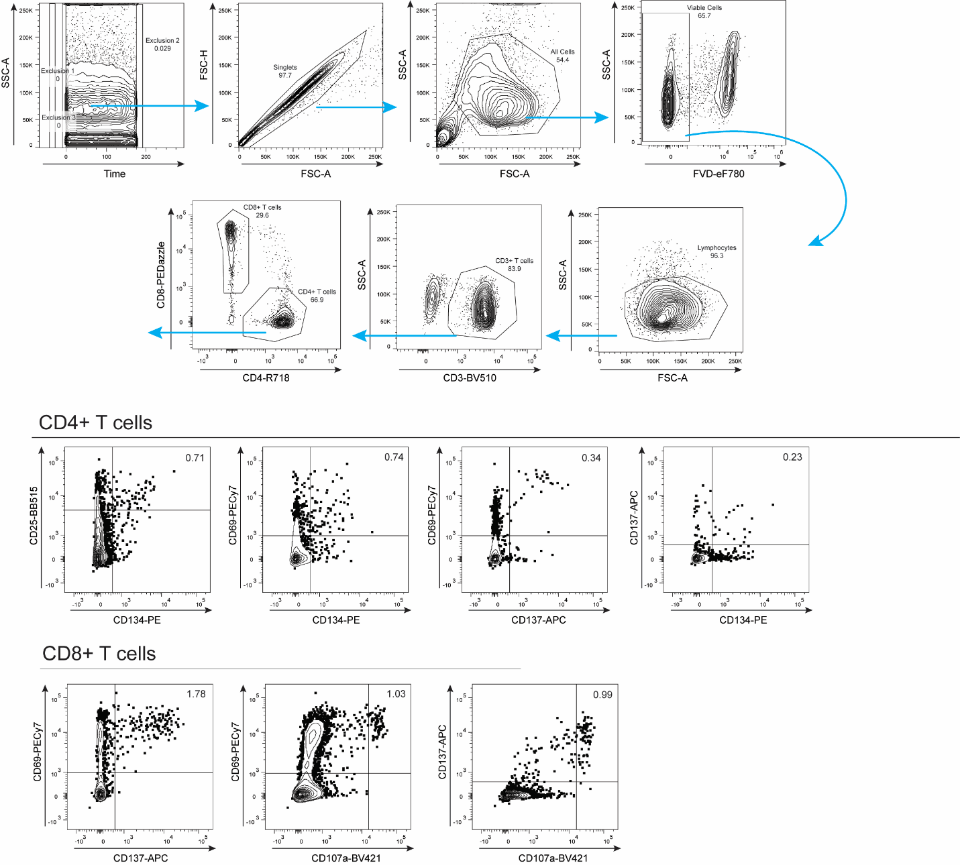

**Figure S11. Flow cytometric gating strategy for multi-centre AIM assay testing.**

Representative flow cytometric gating strategy on CMV-stimulated cryopreserved PBMCs using a simplified panel for multi-centre AIM assay testing. Live CD3^+^ lymphocytes were gated as CD4^+^ or CD8^+^ T cells. AIM^+^ CD4^+^ T cells were defined as CD134^+^/CD25^+^, CD134^+^/CD69^+^, CD137^+^/CD69^+^ or CD134^+^/CD137^+^, and AIM^+^ CD8^+^ T cells were defined as CD137^+^/CD69^+^, CD107a^+^/CD69^+^ or CD107a^+^/CD137^+^. AIM gates were set using the corresponding CytoStim positive control for each donor.

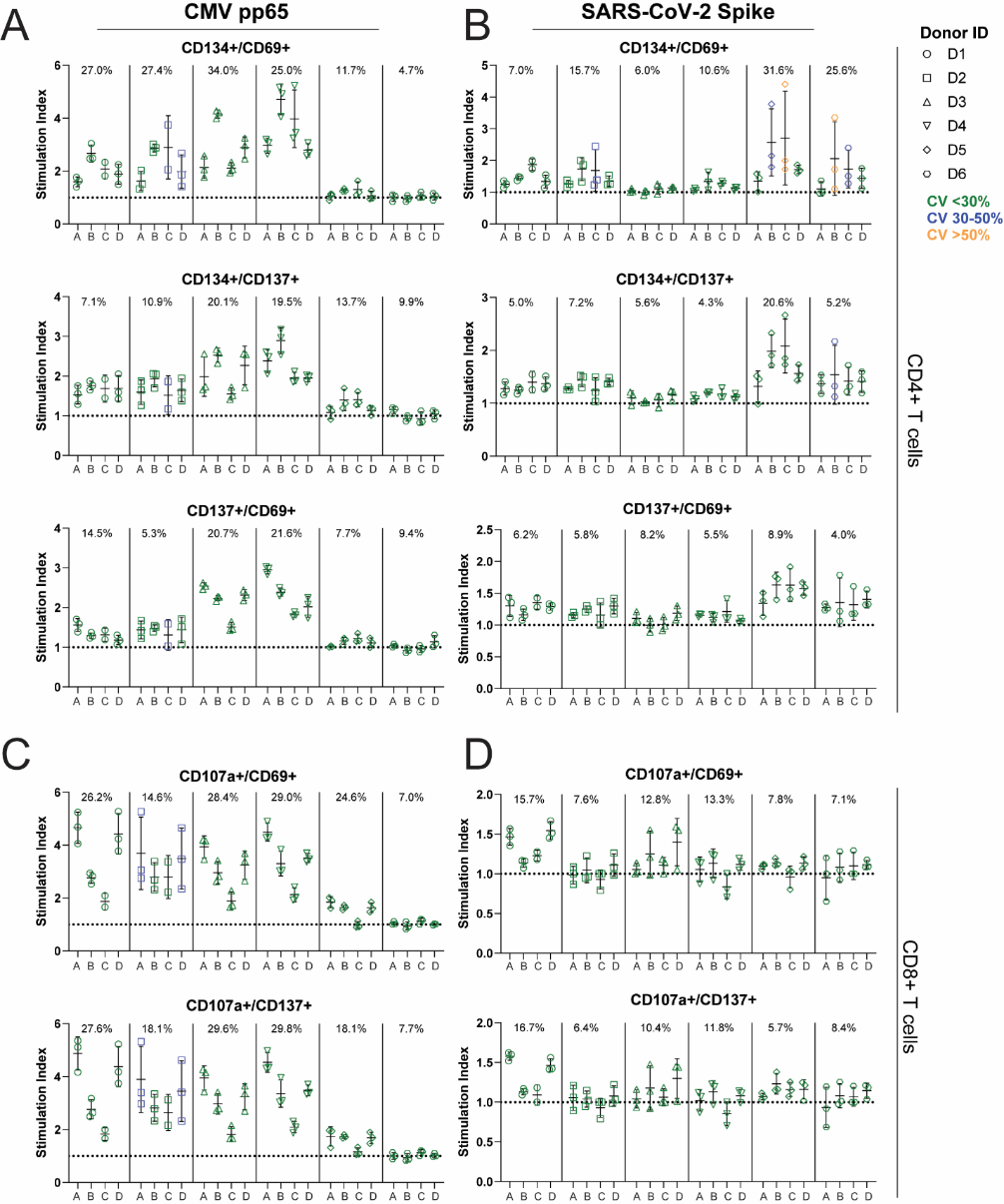

**Figure S12. AIM assays using diverse CD4^+^ and CD8^+^ AIMs are reproducible within and between geographic locations.**

(**A-D**) AIM stimulation indices in CD4^+^ T cells (A and B) and CD8^+^ T cells (C and D) for technical replicate samples of cryopreserved PBMCs stimulated for 20 h with (A and C) CMV pp65 peptides or (B and D) SARS-CoV-2 Spike peptides. Replicates from six donors were assayed in three independent experiments (individual points) at four centres (coded A-D). Figure subpanels separate individual donors. Stimulation index was determined by normalizing pp65 peptide-stimulated AIM^+^ frequencies to the unstimulated control using the Box-Cox correction method described in Figure 2. The means and standard deviations of log_2_-transformed stimulation indices are shown for each AIM. Symbol colours indicate the percent coefficient of variation (CV) within each set of three replicates at each site. The overall CV between sites is shown at the top of the panel for each donor. Related to Figure 6.

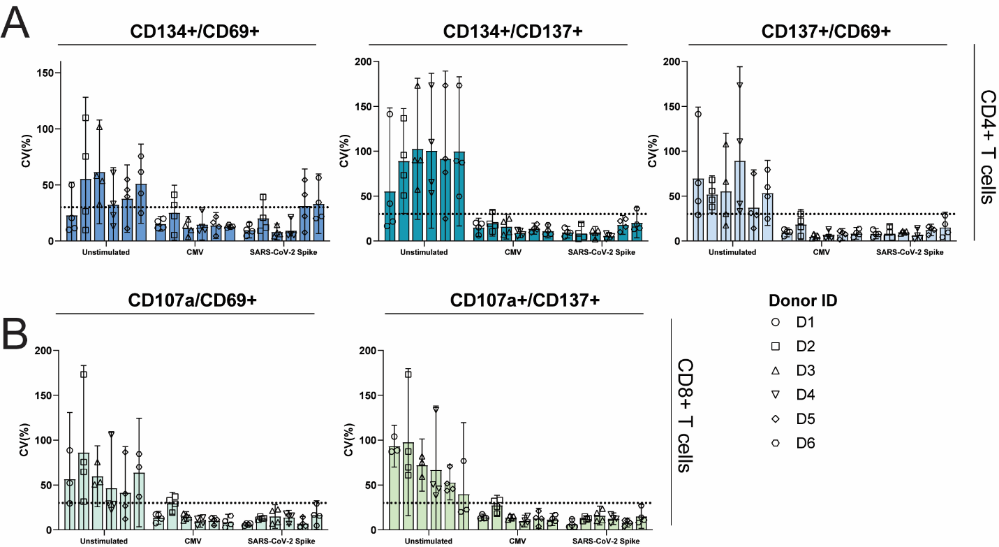

**Figure S13. Intra-site variability in AIM assays differs by AIM pair and stimulant.**

AIM expression was quantified by flow cytometry following a 20-h incubation of technical replicate samples of cryopreserved PBMCs from healthy donors (n = 6) with no stimulant (unstimulated), cytomegalovirus (CMV) pp65 peptides, or SARS-CoV-2 Spike peptides measured at 4 different sites. Stimulation index was determined by normalizing pp65 peptide-stimulated AIM^+^ frequencies to the unstimulated control using the Box-Cox correction method described in Figure 2. Percent CV in (A) CD4^+^ and (**B**) CD8^+^ T cells was calculated across 3 experimental replicates performed across four distinct sites. Means with standard deviation are shown. A percent CV < 30% (dotted line) was assigned as a suitable threshold for reproducibility. Each point represents the CV from three technical replicates within one site; each bar is the mean CV of one donor assayed across the four sites. Related to Figure 6.

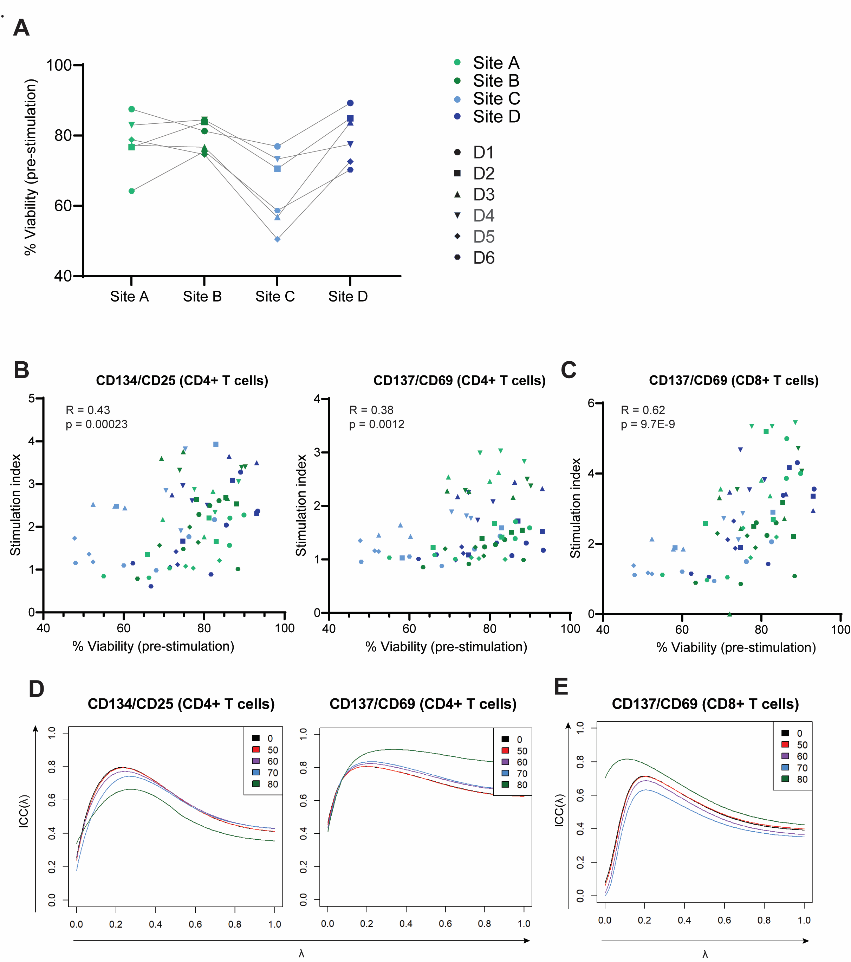

**Figure S14. Effect of viability on AIM stimulation response.**

The percentage of viable cells was analyzed in multi-centre AIM assays on healthy donor PBMCs (n = 6) assayed across 3 experimental replicates at 4 distinct research centres. PBMCs were stimulated with CMV pp65 peptides for 20 h. Viability of each sample was quantified by permeability (propidium iodide or Trypan Blue) staining at the time of antigen stimulation. (**A**) Percentage of viable cells across sites. Each point represents the viability of one donor at the indicated site, as a mean of the three technical replicates within that site. (**B and C**) Correlations between percentage viability and Box-Cox-corrected stimulation index (SI) for (B) CD4^+^ T cell and (C) CD8^+^ T cell AIM responses. Each point represents the viability of one donor at a single site, as a mean of the three technical replicates within that site. (**D and E**) Ratios of biological (between-donor) variance to total variance (intra-class correlation, ICC) plotted as a function of λ using the Box-Cox-corrected SI for (D) CD4^+^ T cell and (E) CD8^+^ T cell AIM responses using different thresholds for excluding samples with low viability. The black line (0%) includes all samples, while the other lines represent ICCs calculated using only samples with at least 50% (red), 60% (purple), 70% (blue) or 80% (green) viability. Related to Figure 6.

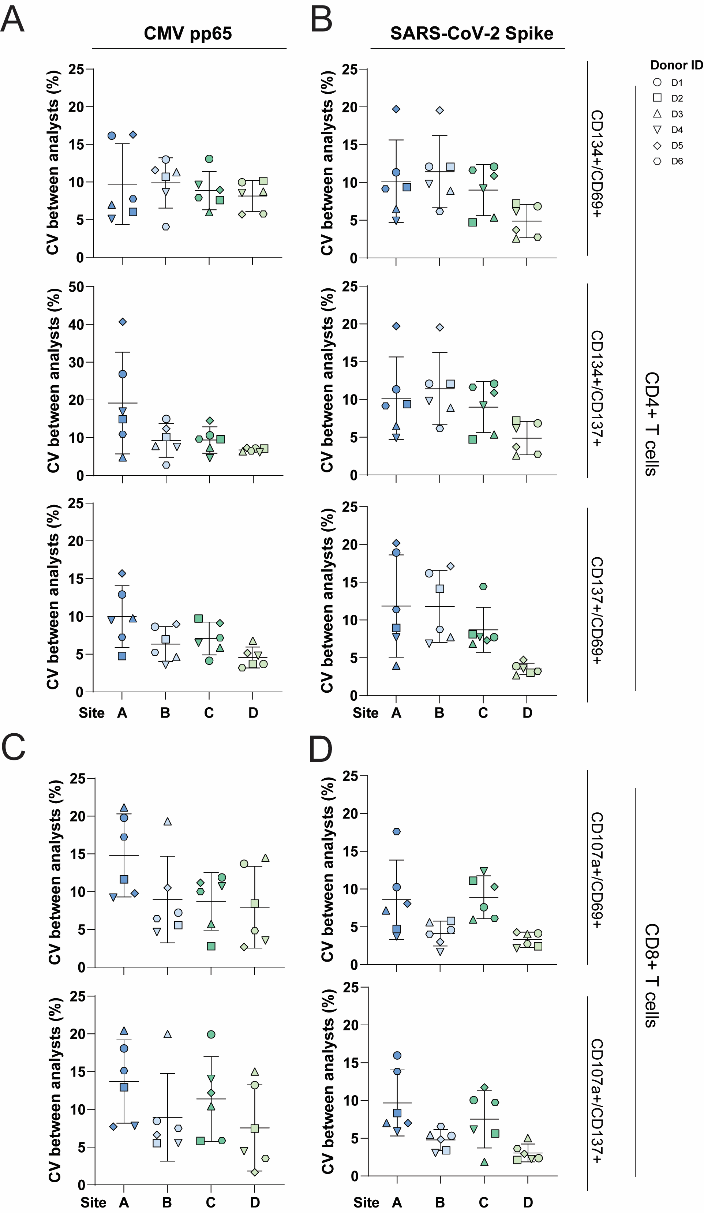

**Figure S15. AIM assays results are subject to variability in manual data analysis.**

(**A-D**) Experienced individuals were assigned identical gating protocols and analyzed the same flow cytometric data files from three technical replicate AIM assays encompassing 20-h stimulation with CMV pp65 peptides (A) or SARS-CoV-2 Spike peptides (B) on healthy donor PBMCs (n = 6) at each of four distinct Canadian research centres (coded A-D for anonymity). The percent CV between individual analysts (n = 3 per site) for analysis of the same dataset is shown for each replicate assay at each of the four sites as an average of CVs from the analysis of each technical replicate. Related to Figure 7.

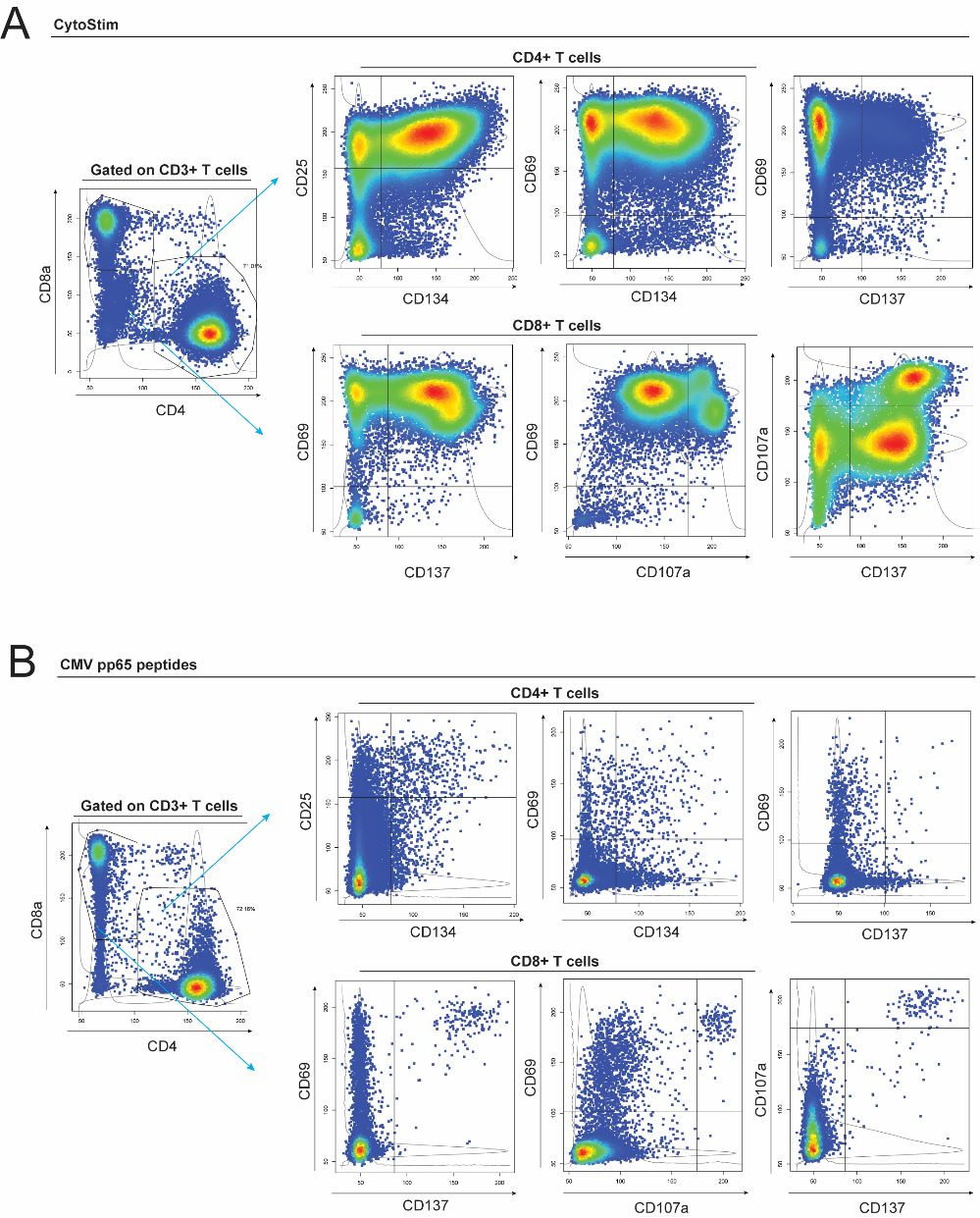

**Figure S16. Representative automated flow cytometric gating strategy for AIM assays**

An automated AIM gating pipeline was created via analysis of AIM assay data from healthy donors (n = 6) assayed in triplicate at each of four research centres. Representative flow cytometric gating approaches are shown for AIM^+^ CD4^+^ and CD8^+^ T cells following a 20-h antigen stimulation of cryopreserved PBMCs. (**A**) A CytoStim-stimulated positive control condition for each donor was used by the automated software to set donor-specific AIM gates based on a nested bivariate segmentation algorithm. (**B**) Gates set by the automated software were subsequently applied to other stimulation conditions for that donor. The CMV pp65 peptide-stimulated condition is shown for example for the same donor as in (A). Related to Figure 7.

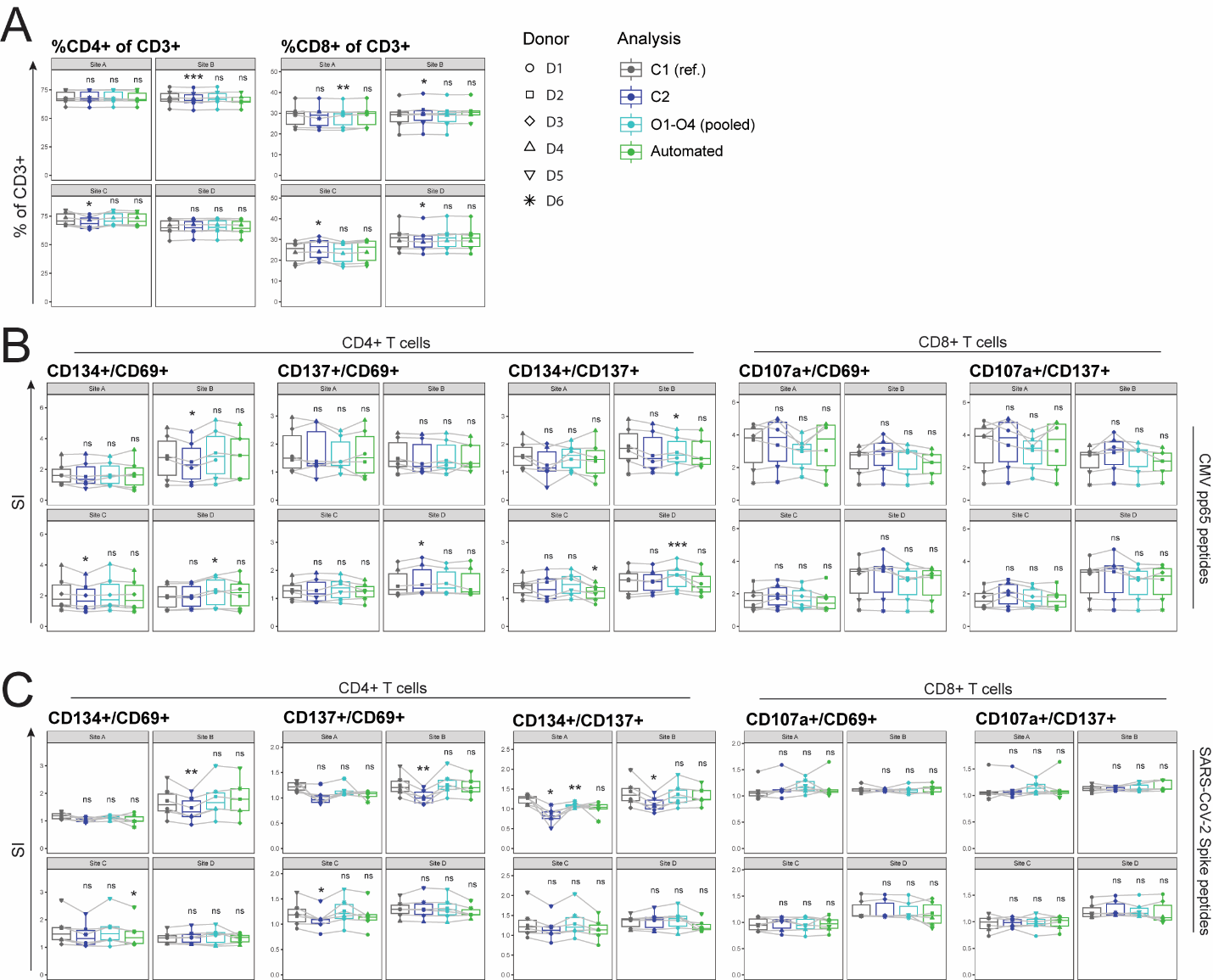

**Figure S17. An automated gating pipeline reliably quantifies AIM responses across a range of CD4^+^ and CD8^+^ AIMs**

AIM assay data from PBMCs (n = 6) assayed in triplicate at four research centres (coded A-D for anonymity) were analyzed manually or via automated gating. A central analyst (C1) defined the gating strategy, provided instructions to the manual analysts (C2 and O1-O4) and oversaw software development for the automated platform, and therefore served as the reference for comparisons. C1, C2 and the automated software each analyzed all data from all sites, while O1-O4 each analyzed data from sites A-D, respectively. (**A-C**) Quantification of (A) bulk CD4^+^ and CD8^+^ T cell populations and (B) CMV- or (C) SARS-CoV-2-stimulated AIM^+^ CD4^+^ and CD8^+^ T cells by manual and automated analysis (reference analyst C1 shown in grey) following a 20-h stimulation. Box-Cox-corrected AIM SI values are shown. P-values represent post-hoc Dunnett’s multiple comparisons test following mixed effects analysis for paired comparisons between each analyst and C1. ns, not significant (p > 0.05); *, p ≤ 0.05; **, p ≤ 0.01; ***, p ≤ 0.001; ****, p ≤ 0.0001. Related to Figure 7.

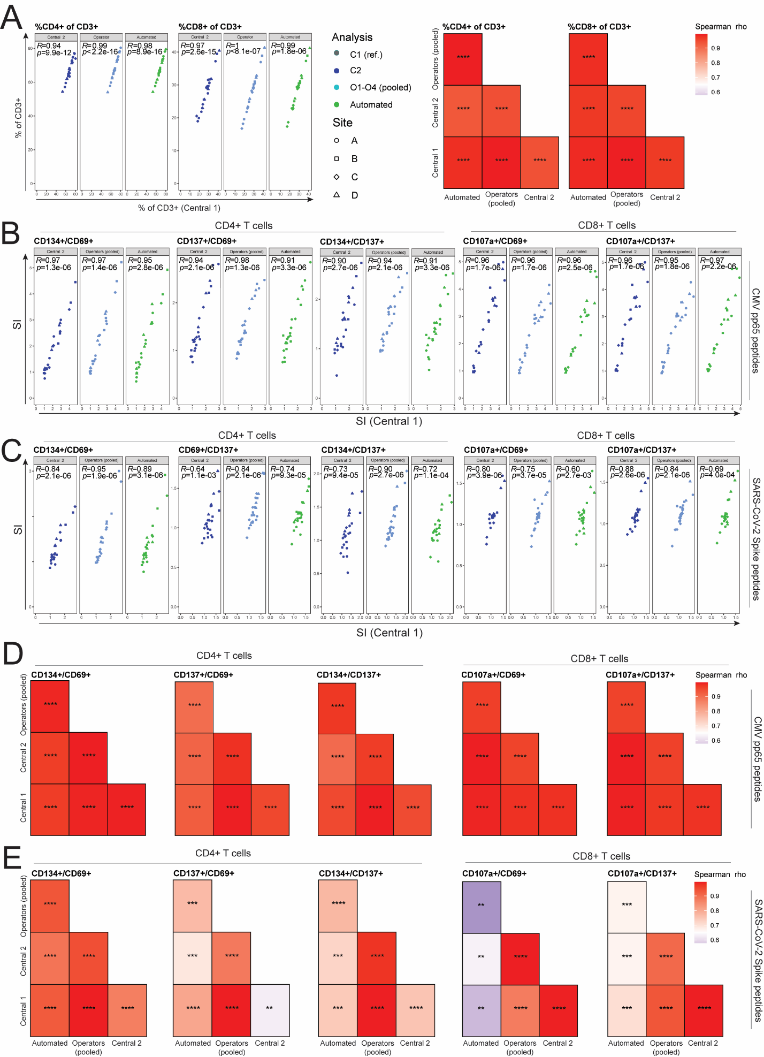

**Figure S18. AIM SI values generated by automated flow cytometric analysis correlate with manually analyzed results across diverse AIM pairs**

AIM assay data from healthy donor PBMCs (n = 6) assayed in triplicate at each of four distinct Canadian research centres (coded A-D for anonymity) were analyzed manually and via automated gating. A central analyst (C1) defined the gating strategy, provided instructions to the manual analysts (C2 and O1-O4) and oversaw software development for the automated platform, and therefore served as the reference for comparisons. C1, C2 and the automated software each analyzed all data from all sites, while O1-O4 each analyzed data from sites A-D, respectively. (**A-E**) Spearman correlations between bulk (A) CD4^+^ and CD8^+^ T cell frequencies (A) and (B and D) CMV- or (C and E) SARS-CoV-2- (C and E) stimulated CD4^+^ and CD8^+^ AIM SI values for manual and automated analysis of multi-centre 20-h AIM assay data. Comparisons are shown against (A-C) the reference analyst and (D and E) for each analysis against all others in a correlation matrix. ns, not significant (p > 0.05); *, p ≤ 0.05; **, p ≤ 0.01; ***, p ≤ 0.001; ****, p ≤ 0.0001. Related to Figure 7.

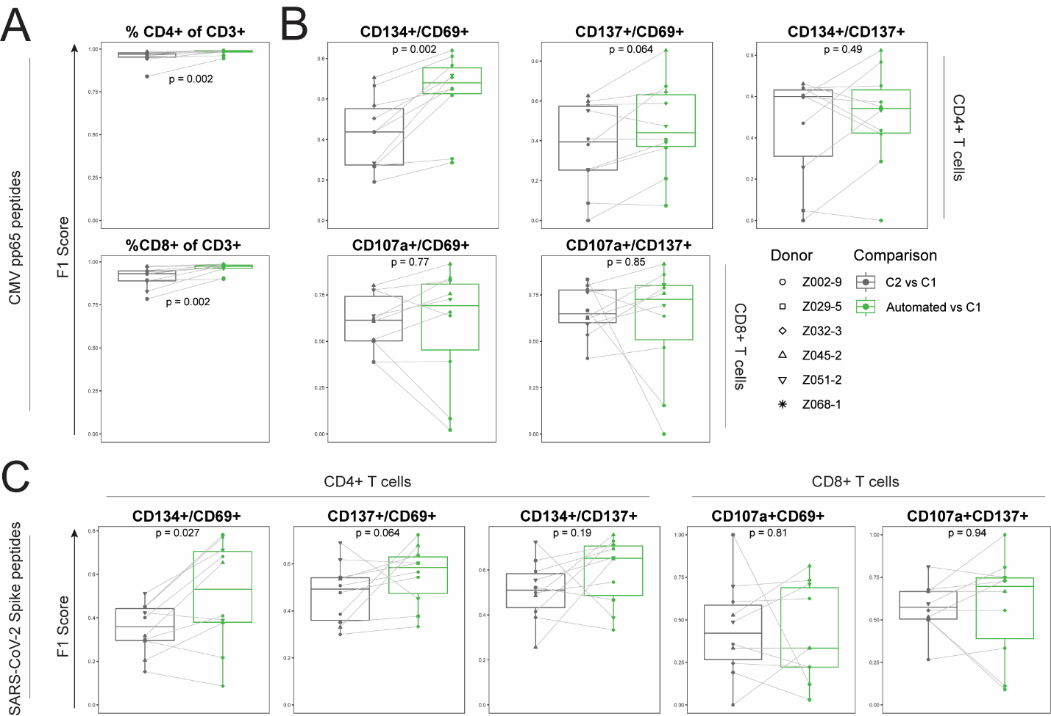

**Figure S19. Similar AIM^+^ populations are identified by automated and manual analysis of AIM assay flow cytometry data**. AIM assay data from healthy donor PBMCs (n = 6) assayed in triplicate at each of four distinct Canadian research centres (coded A-D for anonymity) were analyzed manually and via automated gating. A central analyst (C1) defined the gating strategy, provided instructions to the manual analysts (C2 and O1-O4) and oversaw software development for the automated platform, and therefore served as the reference for comparisons. C1, C2 and the automated software each analyzed all data from all sites, while O1-O4 each analyzed data from sites A-D, respectively. (**A-C**) F1 score comparisons for manual (C2) *vs.* C1 (grey) and automated *vs.* C1 (green) for analysis of bulk CD4^+^ and CD8^+^ T cell populations (A) and AIM^+^ CD4^+^ and CD8^+^ T cells following stimulation with CMV pp65 (B) or SARS-CoV-2 (C) peptides for 20 h. Each point represents the F1 score from a unique donor-site combination, with p-values calculated via paired Wilcoxon signed-rank test after averaging F1 scores from technical replicates. Related to Figure 7.

**Table S1. Extended flow cytometry panel for detection of CD4^+^ Tconv, CD4^+^ Treg^a^ and CD8^+^ T cell AIM responses. Related to Figures 1-5 and S1-S9, and STAR Methods.**

| Antigen | Fluorochrome | Clone | Staining |
| --- | --- | --- | --- |
| CD3 | BUV395 | UCHT1 | Extracellular |
| CD4 | SB600 | SK3 | Intra-stimulation |
| CD8 | BUV496 | RPA-T8 | Extracellular |
| CD25 | BB515 | 2A3 | Extracellular |
| CD274 | BV711 | B7-H1 | Intra-stimulation |
| CD154^b^ | BV785 | 24-31 | Intra-stimulation |
| CD69 | PED | FN50 | Extracellular |
| CD134 | PE | L106 | Extracellular |
| CD137 | APC | 4B4-1 | Extracellular |
| CD107a | BV421 | H4A3 | Intra-stimulation |
| FVD and CD14 | eF780 | 61D3 | Extracellular |
| Helios^a^ | AF700 | 22F6 | Intracellular |
| FOXP3^a^ | PE-Cy7 | 236A/E7 | Intracellular |
| CD39^a^ | BV510 | A1 | Intra-stimulation |

^a^HELIOS, FOXP3 and CD39 were included only when Treg AIM assays were of interest.

^b^An intra-stimulation α-CD40 blocking antibody (1 µg/mL) is included when CD154 staining is desired.

**Table S2. Clinical characteristics of study participants^a^**

|  | Intra-site variability (**Figure 1**)  (n = 5) | Media (**Figure S2**)  (n = 4) | Cell source (**Figures 3 and S6**)  (n = 6) | Time course (**Figures 4 and S7**)  (n = 4) | Treg AIM assay (**Figures 5 and S8**)  (n = 9) | Multi-site testing & automated gating (**Figures 2, 6 and 7)**  (n = 6) |
| --- | --- | --- | --- | --- | --- | --- |
| Age | 46 [32, 48] | 35 [32, 42] | 36 [28, 48] | 29 [24, 34] | 32 [25, 46] | 46 [29, 55] |
| **Sex** | | | | | | |
| Male | 3 [60] | 1 [25] | 2 [33] | 2 [50] | 5 [56] | 3 [50] |
| Female | 2 [40] | 3 [75] | 4 [67] | 2 [50] | 4 [44] | 3 [50] |

^a^Shown are medians with interquartile range (continuous variables) or counts and percentages (categorical variables).

**Table S3. Simplified flow cytometry panel for multi-centre AIM assay testing. Related to Figures 6, 7 and S10-S16, and to the STAR Methods.**

| Antigen | Fluorochrome | Clone | Staining |
| --- | --- | --- | --- |
| CD3 | BV510 | UCHT1 | Extracellular |
| CD4 | R718 | SK3 | Intra-stimulation |
| CD8 | PEDazzle | RPA-T8 | Extracellular |
| CD25 | BB515 | 2A3 | Extracellular |
| CD69 | PE-Cy7 | FN50 | Extracellular |
| CD134 | PE | L106 | Extracellular |
| CD137 | APC | 4B4-1 | Extracellular |
| CD107a | BV421 | H4A3 | Intra-stimulation |
| FVD | eF780 |  | Extracellular |
